# Proteome-wide Prediction of the Functional Impact of Missense Variants with ProteoCast

**DOI:** 10.1101/2025.02.09.637326

**Authors:** Marina Abakarova, Maria Inés Freiberger, Arnaud Liehrmann, Michael Rera, Elodie Laine

**Affiliations:** Sorbonne Université, CNRS, IBPS, Department of Computational, Quantitative and Synthetic Biology (CQSB, UMR7238), 75005 Paris, France; Université Paris Cité, Functional and Adaptive Biology (UMR8251), 75013 Paris, France; Institut Universitaire de France (IUF)

**Author notes:** Corresponding authors: Michael Rera, Elodie Laine.

**Keywords:** Protein mutation, variant effect prediction, developmental lethality, protein function, *Drosophila melanogaster*, functional sites discovery

## Abstract

Dissecting the functional impact of genetic mutations is essential to advancing our understanding of genotype-phenotype relationships and identifying therapeutic targets. Despite progress in sequencing and genome editing technologies, proteome-wide mutation effect prediction remains challenging. Here we show that evolutionary information alone enables accurate prediction of mutation effects across entire proteomes. ProteoCast is a scalable and interpretable computational method that leverages protein sequence conservation to classify genetic variants and identify functionally important protein sites. We apply ProteoCast to the complete *Drosophila melanogaster* proteome (22,000 isoforms, 300 million mutations) and validate it against nearly 400,000 natural and experimental variants. It correctly classifies 85% of known lethal mutations as functionally impactful versus 13-18% of population variants. ProteoCast-guided genome editing experiments confirm these predictions. Moreover, ProteoCast successfully identifies functionally important protein modification sites and binding motifs. ProteoCast provides a publicly available resource and deployable pipeline for studying gene function and mutations in any organism.

## Introduction

More than 50,000 single nucleotide mutations are associated with genetic disorders in humans [1], and correcting them at the genomic level could provide curative treatments for various diseases [2]. The advent of CRISPR (Clustered Regularly Interspaced Short Palindromic Repeats) technology has revolutionised genetics by enabling precise and efficient DNA editing as well as establishing targeted mutational screens as a powerful forward genetic approach to identify novel druggable targets [3]. When combined with high-throughput sequencing, which enables the analysis of inter-individual and inter-tissue genomic variability, these single-point genome editing tools allow unprecedented exploration of the functional significance of genetic variants. They provide effective means for assessing their contributions to disease susceptibility, drug responses and, more generally, to complex traits.

However, the vast number of possible substitutions makes it challenging to select the optimal amino acid target(s) for genome editing. To prioritise single-point genetic modifications for CRISPR experiments, researchers often use conservation scores derived from multiple sequence alignments (MSAs), whole-exome sequencing to identify patient-specific mutations, or statistical tools like Genome-Wide Association Studies (GWAS) to link variants to specific diseases or phenotypes. While invaluable, these methods provide only a limited view of impactful mutations across the entire proteome. Computational predictive models can address these limitations, offering a more comprehensive approach to understanding the functional impact of mutations.

The last decades have witnessed a profusion of protein variant effect predictors. Modern methods generate a mutational landscape - namely, the predicted effects of all possible substitutions at every position within a query protein by leveraging protein sequence data across species. They vary widely in complexity and number of parameters, ecompassing explicit evolutionary models [4–7], family-specific deep learning models [8, 9], and universal protein language models (pLMs) [10–15]. A couple of methods focusing on human applications are equipped with variant classification and uncertainty estimation to facilitate clinical interpretation. Specifically, the EVE method interprets family-specific variational autoencoder score distributions as a mixture of two Gaussians, one representing pathogenic variants and the other benign variants. This approach assumes a direct correspondence between the underlying data distribution and biological variant classes.

Additionally, some predictors are trained on variants with known phenotypes or polymorphisms observed in the population [12, 16]. These population-based predictors, however, do not typically perform better than population-free methods and tend to suffer from circularity issues and pervasive ancestry biases [17]. Furthermore, a new generation of predictors [12, 18–21] systematically integrates high-quality 3D structural models [22]. However, they have shown limited improvement over purely sequence-based methods [23], and the structural contribution is sometimes unclear.

Systematic benchmarking studies [23–25] have demonstrated that predictive performance against deep mutational scans is a good indicator of a variant effect predictor’s ability to identify pathogenic mutations in humans, with modern methods outperforming traditional approaches such as SIFT [4] and PROVEAN [26] at this task. The results highlighted GEMME, a fully interpretable, unsupervised, MSA-based method, as one of the top-performing models [7]. By focusing on how protein residues segregate along the topology of evolutionary trees, GEMME has proven instrumental in studying protein stability, function, and disease mechanisms [27–30]. Notably, GEMME is 100 to 10,000 times faster than deep learning predictors such as ESM-2 [11], EVE [8], and PoET [14], and it does not require high-end GPU resources [7, 12]. When paired with the ColabFold protocol, which provides cost-effective, small, and diverse MSAs, GEMME enables computing mutational landscapes for tens of thousands of proteins within just a few days [31], making it particularly attractive for large-scale studies.

In this study, we expand on GEMME by developing ProteoCast, a scalable fully automated computational framework for facilitating proteome-wide single-point genetic modification prioritisation, and demonstrate its application in *Drosophila melanogaster*. ProteoCast seamlessly estimates the impact of missense mutations on protein function, computes MSA-based confidence metrics, and classifies variants by accounting for the regimes underlying the predicted score distribution (Fig. 1). It determines which residues are sensitive to mutations and maps this information on protein 3D structures. Additionally, it identifies putative binding and regulatory sites in unstructured regions by segmenting the predicted mutational landscape. We validated ProteoCast on a comprehensive benchmark dataset we compiled of over 386,000 single-point mutations from *D. melanogaster*, encompassing genetic variation under natural or artificial selection and known hypomorph and lethal mutations.

**Figure 1.**
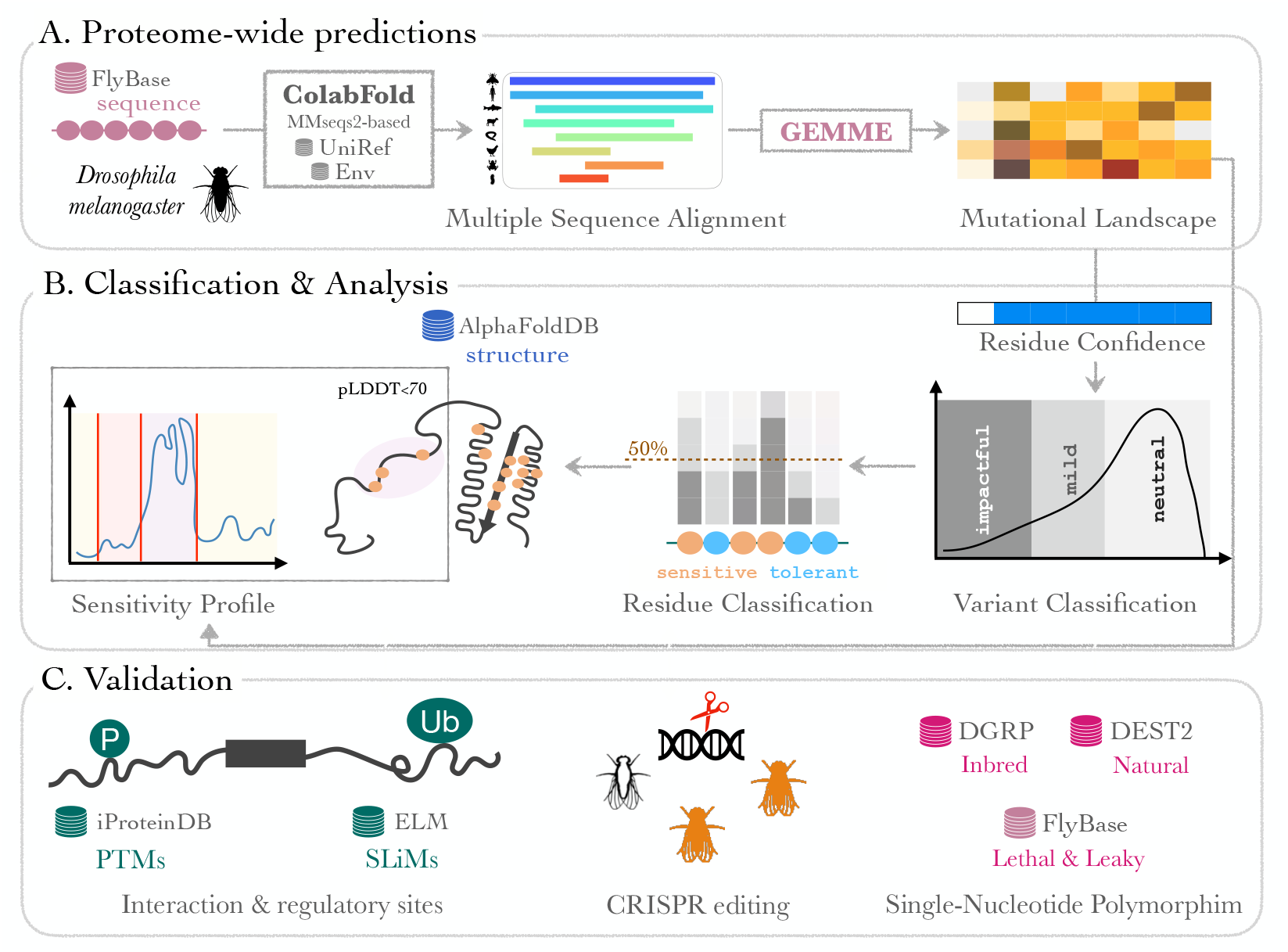
Schematic representation of ProteoCast workflow. A. The workflow starts by generating a Multiple Sequence Alignment for each *Drosophila melanogaster* proteoform sequence using the ColabFold MMseqs2-based pipeline against UniRef and environmental sequence databases. GEMME then produces an Lx20 matrix representing the proteoform complete single-site Mutational Landscape, with L the number of amino acid residues. B. ProteoCast filters out unreliable predictions (Residue Confidence, based on input MSA assessment) and then classifies the remaining variants as impactful, mild or neutral (Variant Classification) by fitting a Gaussian mixture model to the raw score distribution. It then determines whether each residue is sensitive or tolerant to mutations depending on its proportion of impactful variants (Residue Classification). Proteocast further maps per-residue mutational sensitivity (average raw score) to 3D models from AlphaFoldDB. It segments the mutational sensitivity profile to identify potential binding and regulatory sites in unstructured regions. C. ProteoCast validation through CRISPR editing experiments and benchmarking against known post-translational modification (PTM) sites and short linear motifs (SLiMs), as well as single-nucleotide polymorphism, either observed in inbred or wild populations, or associated with deleterious outcome such as developmental lethality or partial loss of function.

Although much genetic research focuses on humans, we selected *D. melanogaster* as our model organism due to its rich and well-documented genetic and phenotypic data accumulated over a century of research. This extensive body of knowledge offers unique insights into natural genetic variability and the functional consequences of mutations. We used three key public resources: the Drosophila Genetic Reference Panel (DGRP) [32], a collection of 205 inbred lines, the community-generated population genomic resource Drosophila Evolution over Space and Time (DEST v2.0) [33], a collection of 530 high-quality pooled libraries from *>*32,000 flies collected across six continents over more than a decade, and FlyBase [34], a comprehensive database on Drosophila genetics that has aggregated the phenotypic effects of point mutations for over 30 years.

With approximately 75% of human genetic diseases being associated with genes conserved in fruit flies, the relevance of Drosophila as a model organism for human disease is clear [35]. Our findings, therefore, are advancing our understanding and ability to intervene in functional genomics, with a broad applicability to human health and diseases. ProteoCast and all generated and analysed data are freely available to the community at https://proteocast.ijm.fr/drosophiladb/. ProteoCast can be run on any protein sequence from any organism using the Docker image at https://hub.docker.com/r/marinaabakarova/proteocast/ or through the webserver at https://proteocast.ijm.fr/.

## Results

We used ProteoCast to generate full single-site mutational landscapes over the entire *D. melanogaster* proteome, including all referenced protein isoforms (proteoforms) resulting from alternative splicing (Fig. 1A, see also Methods). We estimated the functional impact of every possible substitution at every position for 22,169 unique proteoforms, covering 99% of the Drosophila proteome (Table S1). We provide predictions of variant effects for about 293 million missense mutations enriched with detailed information to help guide users in interpreting predictions more effectively. These metadata include confidence scores, variant functional classification as neutral, mild, or impactful, as well as each residue’s sensitivity to mutations. Our online database allows for querying any Drosophila proteoform and interactively exploring the associated ProteoCast predictions. Below, we provide a graphical summary of the prediction pipeline (Fig. 1) and functional validation of the predicted functional impacts.

### Sufficient evolutionary information for most of the variants

To help users assess the reliability of predictions based on input data quality, ProteoCast provides confidence metrics that flag cases, proteins or residues, where insufficient evolutionary information may compromise prediction accuracy (Fig. 1B, *Residue Confidence*). At the protein level, we consider that mutational landscapes predicted from less than a couple hundred sequences may be of low quality. This cutoff is based on our previous findings [31] and is calibrated for the MMseqs2-based MSA generation protocol implemented in ColabFold [36, 37] that maximises diversity while minimising the number of sequences.

At the residue level, ProteoCast uses the predicted scores’ dispersion over the 19 possible substitutions as a proxy for the input alignment information content. A small dispersion indicates that ProteoCast is not able to distinguish substitutions from one another. On the one hand, this situation typically occurs upon strong selection pressure, when a position is highly conserved in evolution and thus does not tolerate any substitution. A high conservation level implies a limited subset of amino acids observed in the input MSA and a low gap content. On the other hand, a narrow distribution of predicted scores may reflect lack of evidence from the input MSA, for instance in the case of highly gapped or highly divergent positions. In this situation, the conservation level will be low (Fig. S1). Hence, ProteoCast considers that positions with a small score dispersion, a low conservation level, and a low coverage or diversity in the input MSA lack supporting evidence and flags them as unreliable (see Methods for a more formal definition and Fig. S1, insert).

Over the whole *D. melanogaster* proteome, ProteoCast filtered out about 12% of all unique proteoforms according to the global (per-protein) criterion (Fig. S1). It further detected unreliable predictions for *∼* 460,000 residues (*∼*3%) in the remaining proteoforms via the local (per-residue) confidence metric (Fig. S1). Overall, the predictions labelled as unreliable based on the global or local confidence metrics concerned less than 10% of all residues Moreover, for most of the proteoforms, more than 70% of all possible amino acid substitutions (19 multiplying the protein length) are in fact observed in the input alignment, ensuring high-resolution predictions. These results underscore the quality and abundance of the evolutionary information available for the fly proteome.

### High correlation between predicted effects and organismal fitness

ProteoCast predicts a protein’s complete single mutational landscape based solely on evolutionary information specific to that protein, without considering its biological function nor interactions with other proteins. To evaluate the relevance of these protein-centered predictions for organismal fitness, we created a custom benchmark of missense mutations resulting from single nucleotide polymorphisms (Fig. 1C, *Single-Nucleotide Polymorphism*). This benchmark comprises all inbred missense DGRP polymorphisms, all natural missense DEST2 polymorphism as well as all ethyl methanesulfonate (EMS) point mutations referenced in Flybase. We used Flybase’s experimentally supported annotations labeled as hypomorph - indicating reduced gene function - or as developmentally lethal in flies (Table S2, Fig. S2). The benchmark is highly imbalanced, comprising approximately *∼*137,000 DGRP and *∼*270,000 DEST2 missense mutations spread over more than 10,000 genes, *∼*1,000 lethal mutations over almost 500 genes, and 148 hypomorph (Fig. S2B).

We illustrate ProteoCast analysis with the Yorkie protein (Fig. 2A-B), a key component of the Hippo pathway involved in apoptosis and cell proliferation, among other developmental processes [38]. We highlight four mutations on the landscape predicted by ProteoCast in Fig. 2A: the lethal mutations P65L and Y259N rank among the top 5% most negative predictions (dark cells), clearly distinguishable from the naturally occurring DEST2 variants N169T and Q177P (light cells). We map all annotations on the predicted score distribution in Fig. 2B, where mutations inducing lethality (in red) are shifted toward lower (more negative) scores compared to population variants (in blue).

**Figure 2.**
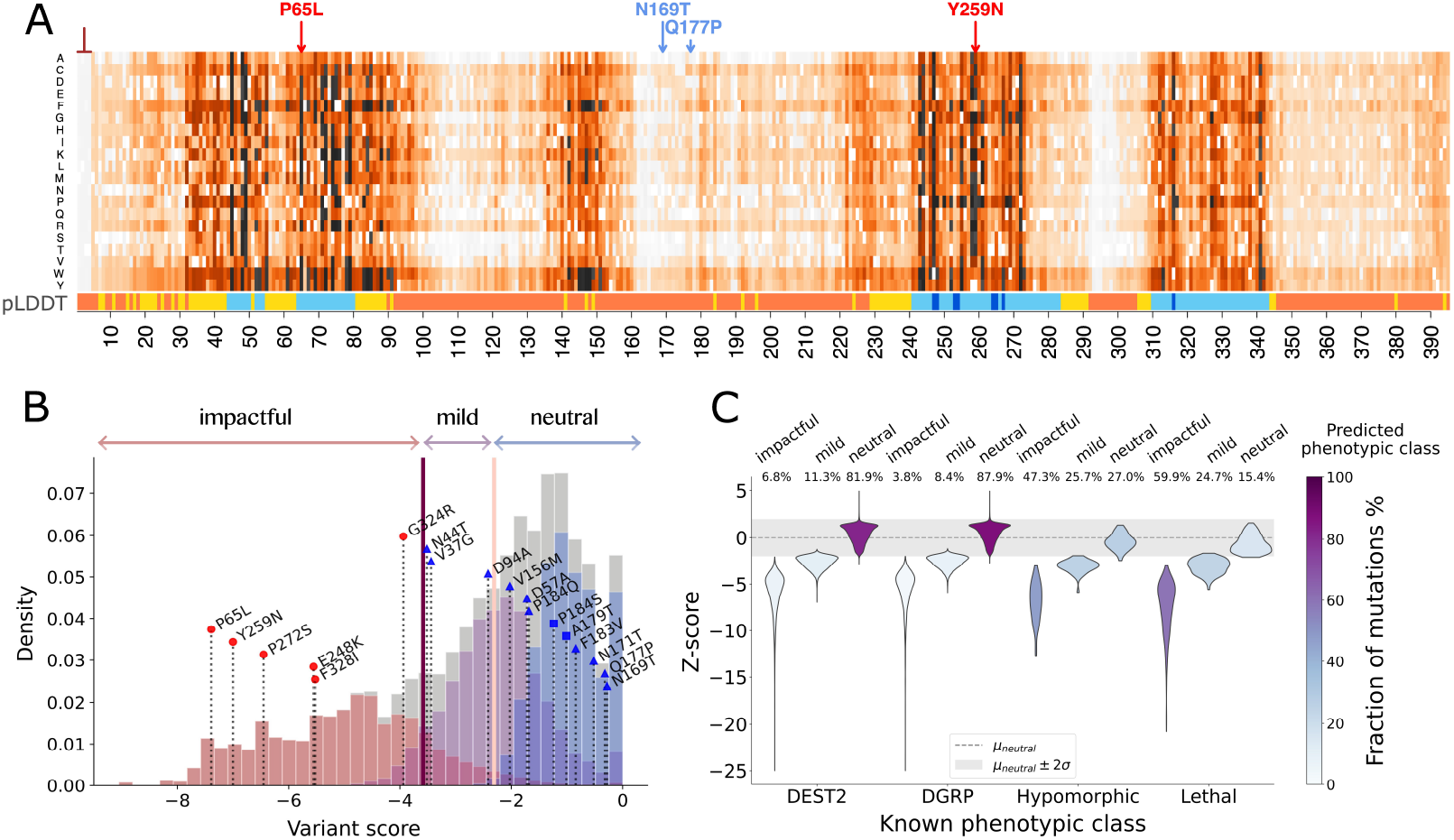
Variant classification using ProteoCast. **A.** Full single-mutational landscape computed for Yorkie protein (*yki* gene, Flybase proteoform id: FBpp0288697), a key downstream effector in the Hippo/SWH (Sav/Wts/Hpo) signaling pathway. Each cell of the matrix contains a numerical estimate of the functional impact of an amino acid substitution (y-axis) at a given residue position (x-axis) on the protein function. Darker colours indicate stronger effects, and white cells correspond to the wild-type amino acids. The inverted T-shaped symbol at the N-terminus marks low-confidence predictions (Residue Confidence). The highlighted mutations provoke developmental lethality – as referenced in FlyBase (P65L, and Y259N, in red) or occur in DEST2 (N169T, and Q177P, in blue). The pLDDT (predicted local distance difference test) score, shown as a colour bar below the matrix, provides a confidence measure of the AlphaFold2-predicted 3D structure, orange: very low, yellow: low, light blue: high, dark blue: very high. Residues 241–274 and 310–343 correspond to the well-characterised and well-structured WW domains. B. ProteoCast classification of Yorkie mutations where the pale pink and deep purple vertical lines indicate the neutral/mild and mild/impactful frontiers, respectively. All 18 known mutations in our benchmark are indicated with red dots for lethal, blue triangles for DEST2, and blue squares for DGRP. C. Distributions of Z-scores for DGRP and DEST2 polymorphisms, as well as hypomorph and lethal mutations annotated in FlyBase, depending on whether they are classified as impactful, mild or neutral. We computed the Z-scores with respect to the distribution of estimated effects for the neutral class within each protein. Three mutations with scores below -25 were excluded from the plot for graphical representation clarity. Source data are provided as a Source Data file.

To systematically and automatically segregate variants, ProteoCast applies a thresholding strategy tailored to each protein by fitting a mixture of Gaussians to the predicted raw score distributions (Fig. 1B, *Variant Classification*, Fig. 2B). We set the number of components to three as it represented an optimal choice for maximising likelihood while avoiding overfitting and overparametrisation across 20 trials of 100 randomly selected proteoforms (Fig. S3A-B, and see Methods). We take the mean of the central component as a threshold to distinguish between *neutral* mutations (scores towards 0) and mutations altering the protein function (lower, more negative scores, Fig. 2B). The latter span a substantially wider range of scores than the former (Fig. S3C-D), suggesting varying degrees of evolutionary constraints against them. To reflect this richness in the evolutionary data, we define two subgroups of variants, namely *mild* and *impactful*, using the intersection between the left-most and central Gaussians (Fig. 2B). The three variant classes exhibit overlapping predicted score distributions with increasing means, from impactful through mild to neutral (Fig. S4).

This classification procedure exhibited high discriminative capability over our benchmark. On the one hand, ProteoCast detected 85% of the lethal mutations and 73% of the hypomorph ones as impactful or mild. On the other hand, it identified 88% of the DGRP variants and 82% of the DEST2 ones as neutral (Fig. 2C and Table S3). The difference between the two datasets may reflect the stronger selection over flies inbred in the laboratory compared to natural variants in living flies. Results obtained for individual proteins are in line with these overall statistics: ProteoCast correctly classified over 80% of the lethal, DGRP and DEST2 mutations in 78%, 82%, and 66% of the cases, respectively (Fig. S5). To further examine the internal structure of our predicted classes, we computed Z-scores measuring how each variant’s raw score deviates from the neutral class distribution (Fig. 2C). Lethal and hypomorph mutations consistently exhibited more negative Z-scores than population-based variants, even within the same predicted class, revealing a continuous gradation of predicted severity that aligns with experimental annotations. Additionally, population polymorphisms with lower Minor Allele Frequency (MAF) tend to be assigned more negative raw scores and Z-scores (Fig. S6A).

By adapting to the shape of each protein’s score distribution, ProteoCast variant classification strikes a better balance between detecting lethal mutations as impactful or mild (recall) and assigning neutral impact on population variants (specificity) than universal thresholds (Fig. S6B-C, Table S3, and Fig. S7). This advantage is especially visible for proteins with near-complete evolutionary sampling of substitutions (*>*95% actually observed in the input MSA), where the predicted scores are shifted toward higher values (Fig. S8A-B). For instance, applying Yorkie’s thresholds to the serine/threonine phosphatase Flapwing (Flw), essential for larval development, would make its two lethal mutations (Y264F and D222N, scores of -1.38 and -0.97) appear as neutral. Thanks to its adaptive strategy, ProteoCast identified these mutations as mild and correctly distinguished them from the DGRP mutation E314D, classified as neutral (Fig. S8C-D).

Taken together, these results show that variant effects inferred from the evolutionary history of individual proteins correlate well with experimentally characterised organismal fitness. The vast majority of mutations observed in viable strains, such as the DGRP lines and DEST2 pooled libraries, are not strongly selected against in evolution. Conversely, higher severity tends to be associated with higher evolutionary pressure detectable by our approach. The consistency between overall performance metrics and individual protein results further demonstrates that our approach generalises well across the entire proteome.

The distinction between mild and impactful categories was not systematically evaluated as available experimental annotations are predominantly qualitative and heterogeneous rather than providing standardised severity gradations. Moreover, the distinction between lethal and hypomorph mutations in our benchmark may be confounded by experimental conditions, as some hypomorph mutations have only been characterised in heterozygous states or under permissive conditions, and could actually be lethal. This complexity explains why the correspondence between experimental categories (lethal/hypomorph) and our predicted classes (impactful/mild) is not straightforward. Nevertheless, our graduated classification captures meaningful severity gradations, as illustrated by insulin receptor mutations where the mild variants R1467C and G1599R cause moderate phenotype (reduced head size) while the impactful variant G1539E produces severe effects (strongly reduced head size, putative null allele) [39].

### ProteoCast effective in guiding genome editing strategies

We used ProteoCast predictions to prioritise variants for experimental genome editing, introducing single-point missense mutations in *D. melanogaster* (Fig. 1C, *CRISPR editing*). First, we selected nine candidate genes with high-quality input alignments, consisting of 10,000 to 30,000 sequences each. These genes were chosen based on the absence of lethal missense mutations reported in FlyBase and a broad range of predicted raw scores, including very negative values (below -8), indicating strong effects (see Methods). We first tested the developmental lethality of RNAi-induced knockdowns for these genes using transgenics under control of the ubiquitous *daughterless::GeneSwitch* inducible driver. The RNAi-mediated knockdown of Naprt (CG3714), which encodes nicotinate phosphoribosyltransferase, an enzyme involved in NAD biosynthesis, resulted in 100% developmental lethality, identifying it as an essential gene during development (Fig. S9).

These results led us to prioritise *Naprt* for genome editing. We selected five variants based on ProteoCast predictions computed for Naprt-PH (662 residues), the longest proteoform crossreferenced in UniProt (Table 1). We first chose the two variants identified by ProteoCast as the most impactful, namely G497P and K532G. Both are located near the putative active sites, as inferred by homology transfer (Fig. 3 and Table S4). We chose the third target residue, E147, among the 25 most sensitive to mutations, as estimated by averaging scores over the 19 possible substitutions (Table 1). The 3D structure of Naprt’s human homolog suggests it could be involved in the protein’s homo-dimeric interface (Fig. 3D and Fig. S10). We substituted it with the amino acid predicted to have the strongest effect, namely Valine (V).

**Table 1.** Residues selected for CRISPR introduction of single amino acid missense mutations. For each selected variant, we report its class (among impactful, mild and neutral), its raw predicted score and associated overall and relative ranks. In addition, we report the mutational sensitivity class (either tolerant or sensitive) and rank of the corresponding wild-type residue. For comparison, we indicate SIFT predicted normalised probabilities for the selected variants. ^*∗*^Residues with pLDDT*<*70 are excluded from the ranking. This leaves us with 533 residues and 10,127 possible variants. The arrows indicate how the values are ordered to compute the ranks. For instance, a variant ranked 1 with an up arrow has the lowest (most negative) raw score. When some values are equal, they are assigned the same (minimum) rank. Sequencing validations of the five mutants are presented in Annex 1.

| Naprt Variant | Variant class | Raw score | Rank overall* | Rank at residue | Residue class | Residue rank* | SIFT scores |
| --- | --- | --- | --- | --- | --- | --- | --- |
| G497P | Impactful | -8.75 | 1 ↑ | 1 ↑ | Sensitive | 1 ↑ | 0.00 |
| K532G | Impactful | -8.73 | 2 ↑ | 1 ↑ | Sensitive | 11 ↑ | 0.00 |
| E147V | Impactful | -8.33 | 52 ↑ | 1 ↑ | Sensitive | 21 ↑ | 0.00 |
| I618P | Neutral | -1.54 | 3,322 ↓ | 1 ↑ | Tolerant | 5 ↓ | 0.21 |
| A262F | Neutral | -0.84 | 1,670 ↓ | 3 ↑ | Tolerant | 2 ↓ | 0.05 |

**Figure 3.**
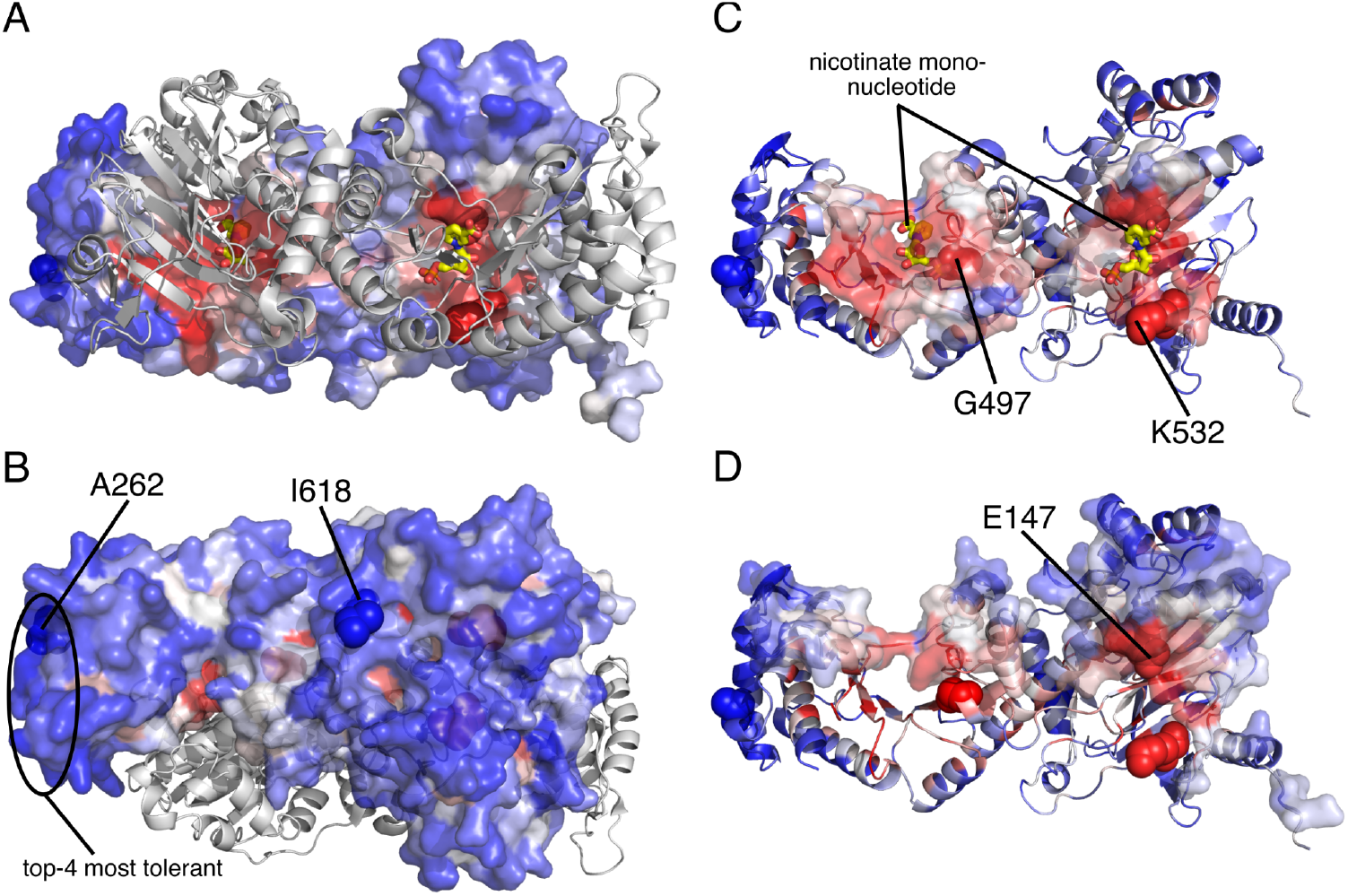
3D mapping of Naprt mutational sensitivity and selected variants. The AlphaFold 3D model of the Naprt-PH proteoform (FlyBase id: FBpp0306840, 662 residues) is colored according to ProteoCast- predicted mutational sensitivity, from blue (low sensitivity) through white to red (high sensitivity). The unstructured regions 1-21 and 301-407, with pLDDT*<*70, are omitted for clarity. The five variants selected for CRISPR validation are highlighted in spheres and labelled. A-B. Complex obtained by superimposing the biologically relevant homo-dimeric arrangement of the human Naprt (PDB id: 4YUB, see also Table S4 and Fig. S10). The two yellow compounds, nicotinate mononucleotides, indicate the putative location of the active sites. They were positioned by superimposing the structure of TaNAPRTase from *Thermoplasma acidophilum* (PDB id: 1YTK). A. Front view. B. Back view. C-D. Transparent surfaces indicate regions potentially involved in binding the substrate or in forming the homo-dimer. C. Residues within 10 Å from the nicotinate mononucleotides. D. Residues within 5 Å from the superimposed human Naprt’s second monomer.

As negative controls, we looked for residues predicted to be tolerant to mutations, with more than 10 out of the 19 possible substitutions predicted as neutral (Fig. 1B, *Residue Classification*). To ensure a rigorous test of ProteoCast’s predictions, we excluded the N-terminal extremity (residues 1-21) and an inter-domain linker (residues 301-407). These regions are poorly conserved and unstructured, and they comprise the vast majority (17 out of 20) of the naturally occurring missense mutations referenced in DGRP and DEST2. We focused on the five most tolerant residues in the structured protein fraction. The top-4 residues form a cluster in 3D space (Fig. 3B), from which we randomly selected a representative residue, A262.

Additionally, we retained I618, the fifth-ranked residue, located in a solvent-exposed helix. To challenge the robustness of ProteoCast’s neutral predictions, we substituted A262 and I618 with amino acids that exhibit starkly contrasting physico-chemical properties (Table 1). Specifically, we mutated the small A262 into Phenylalanine (F), a large aromatic amino acid, and I618 into Proline (P), which has the tendency to break secondary structures. This design allowed us to test whether ProteoCast could correctly predict neutral effects despite substantial changes in amino acid properties.

The single point mutation genome editing was conducted in collaboration with the company WellGenetics following the designs presented in Annex 1 in a *w*^1118^genetic background. The obtained results were perfectly in line with our predictions. Specifically, the introduction of either of the mutations G497P, K532G or E147V did not allow for homozygotization, while introducing I618P or A262F mutations led to viable homozygotes.

To further demonstrate the added value of ProteoCast, we compared it with the popular tool SIFT [40] (Fig. S11). Predictions from the two methods showed overall good agreement (Fig. S11A,C). However, SIFT assigned a score of 0.05 to the variant A262F (Table 1). While this value falls at the boundary of SIFT’s deleterious threshold, both ProteoCast and the CRISPR experiment clearly indicate A262F is neutral. Moreover, ProteoCast offers much finer resolution and higher dynamic range of the mutational effects (Fig. S11B,D). For instance, SIFT assigns the minimal score of 0 to 2,382 variants, representing about a quarter of all possible variants. Among these, ProteoCast classifies 188 variants as neutral, demonstrating its ability to discriminate within what SIFT considers uniformly deleterious. This ability of ProteoCast to discriminate between substitutions across protein sites and also at each protein site constitutes a significant advantage for experimental design and clinical prioritisation.

### Discovery of interaction and regulatory sites in unstructured regions

ProteoCast maps predicted variant scores to proteoform 3D models from the AlphaFold database [22, 41], leveraging its extensive protein structural data. We observed that when GEMME faces difficulty making accurate predictions due to limited evolutionary information, AlphaFold2 similarly struggles to produce reliable 3D models. Specifically, 96% of the residues flagged by ProteoCast as having unreliable predictions are located within protein regions with low- or very low-confidence AlphaFold2-predicted 3D coordinates (predicted Local Distance Difference Test scores below 70, Fig. S12A). Conversely, however, ProteoCast estimated that 94% of the residues with pLDDT scores below 70 had confident variant predictions (Fig. S12B). Hence, AlphaFold2’s inability to model 3D structures does not imply that the evolutionary signal is too weak for ProteoCast to predict variant effects.

Yorkie’s mutational landscape exhibits several local patterns of strong predicted variant effects within low-pLDDT regions (Fig. 2A). While UniProt does not provide any information about these regions, beyond intrinsic disorder detected by MobiDB-lite [42], the high mutational sensitivity of segments 32-98 and 135-160 suggests that they are important for the protein function. To validate this hypothesis, we analysed post-translational modification (PTM) site annotations from iProteinDB [43] and experimental structural data from the Protein Data Bank (PDB) [44]. Both segments exhibit a high concentration of known phosphorylation sites – 10 out of 20 annotated (Fig. S13A), and segment 32-98 corresponds to the TEAD binding domain of Yorkie’s human homolog, YAP1 [45]. This domain is natively unfolded and conditionally adopts a stable conformation upon binding to its partner that is closely mirrored by Yorkie’s AlphaFold2-predicted model (Fig. S13A). Hence, the evolutionary signals detected without any supervision by ProteoCast point at unstructured segments directly involved in protein regulation and interactions.

To systematically and automatically detect such sites, ProteoCast models mutational sensitivity as a piecewise constant continuous parameter and detects both the number of changepoints and their location using the FPOP algorithm [46, 47] (Fig. 1B, *Sensitivity Profile*, and see Methods). The pLDDT profile is used to tune granularity and avoid over-segmenting well-structured regions. For ease of interpretability, the detected segments are colour-coded (Fig. 4A-B and Fig. S13B) depending on whether they have elevated sensitivity compared to both neighbouring segments (purple), only one (red) or none (white). We quantitatively assessed this segmentation over the entire fly proteome, focusing on regions poorly modelled by AlphaFold2 (Fig. 1C, *Interaction & regulatory sites*). Our evaluation benchmark comprises 53,737 PTM sites annotated in iProteinDB database [43] and 72 short linear motifs (SLiMs) from the ELM resource [48]. ProteoCast successfully identified 63% of the PTM sites in segments showing elevated mutational sensitivity (Table S5 and Fig. 4C). This estimate is in line with previous studies reporting between 35% and 65% of phosphosites under evolutionary constraints across eukaryotic proteomes [49, 50]. Regarding SLiMs, 85% partially overlapped with the segments detected by ProteoCast and 61% are fully included in them (Table S5 and Fig. 4C). Overall, these results confirm that SLiMs, and to a lesser extent PTMs, are under selective pressure. Moreover, the strategy of defining segments whose mutational sensitivity is homogeneous and elevated compared to their surrounding context yields a higher detection rate than relying on the absolute mutational sensitivities computed for individual residues (Table S5 and Fig. S14).

**Figure 4.**
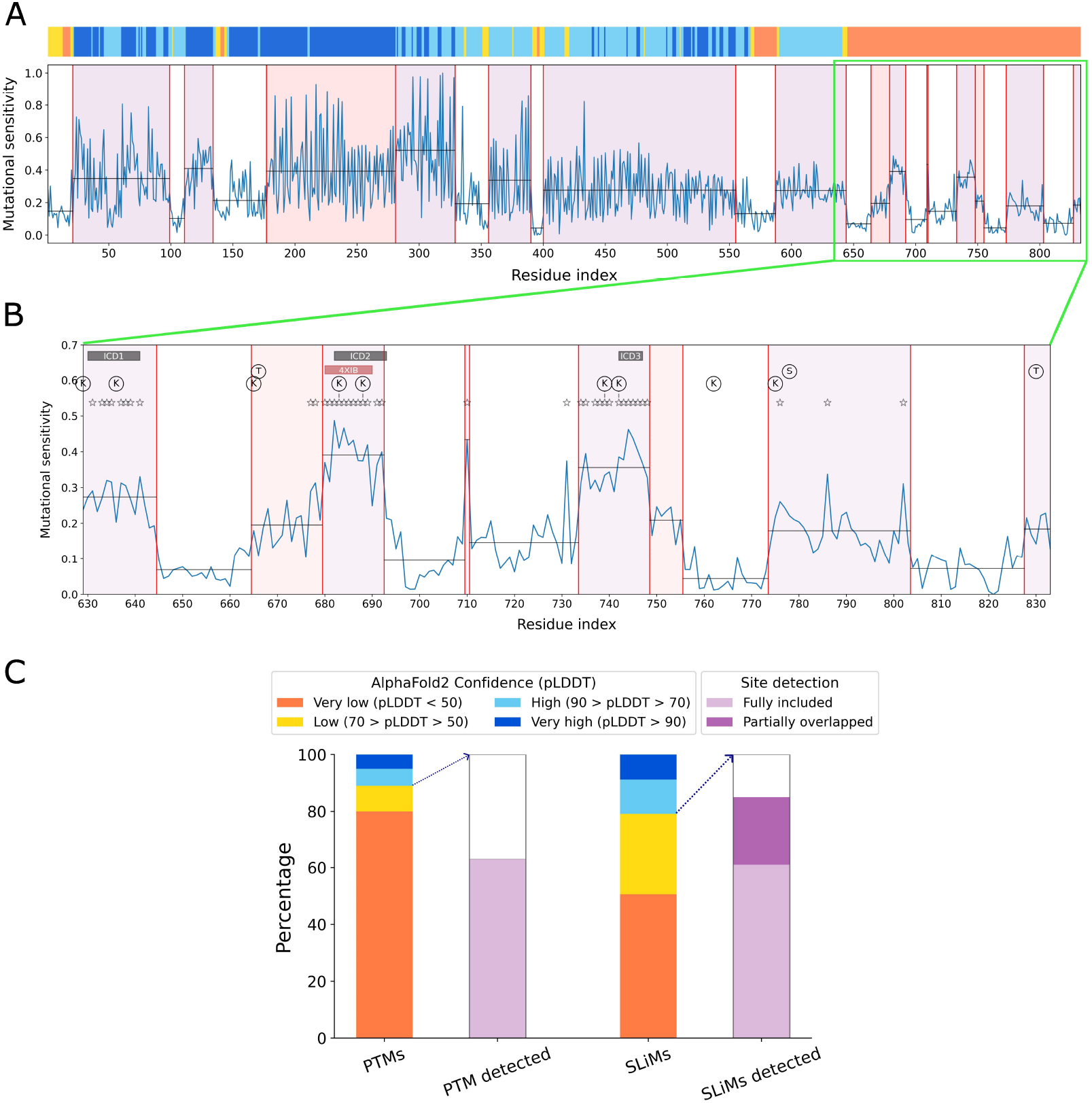
Structural confidence and mutational sensitivity of the Delta protein. A. Top. The pLDDT confidence score from the AlphaFold2-predicted 3D structure is shown as a color bar. Orange and yellow indicate very low to low confidence, while light and dark blue represent high to very high confidence regions. Bottom. Mutational sensitivity as estimated by the average of the scores predicted for the 19 possible substitutions. The curve is segmented using the FPOP algorithm (see Methods). The identified segments are colored red if their mean value (horizontal line) is higher than one neighbouring segment, and purple if it is higher than both neighboring segments. B. Close-up view of ProteoCast sensitivity profile for the C-terminal region from residues 629 to 833, where the stars highlight residues sensitive to mutations, according to ProteoCast classification. The black boxes mark experimentally validated binding motifs for ubiquitylation by Mindbomb1 and Neuralized [51]. The pink box indicates the region whose structure was experimentally resolved in complex with Mind bomb 1 (MIB1) (PDB code: 4XIB, residues 680 to 690). Ubiquitination sites are indicated by circled “K,” and phosphorylations by circled “S” and “T”. C. ProteoCast detection of PTM sites and SLiMs over the *D. melanogaster* proteome. The left bars indicate the proportion of known PTM sites and SLiMs in the different AlphaFold pLDDT bins. The right bars give the proportions of PTM sites and SLiMs partially overlapping with or fully included in segments detected by ProteoCast as having elevated sensitivity. Source data are provided as a Source Data file.

We further used ProteoCast profiling to decrypt the sequence of the unstructured C-terminal part of the Delta protein (833 residues), a ligand of the Notch signalling pathway (Fig. 4A-B). The residues predicted as sensitive to mutations (*>*50% substitutions classified as neutral, indicated as stars) are mostly clustered in three segments identified by ProteoCast as having higher sensitivity than their background context (in purple). While UniProt does not provide any functional information about these segments, we found evidence in the literature for their direct implication in regulating Delta’s ubiquitination and activation by Mindbomb1 and Neuralized (Fig. 4B, black and pink boxes) [51]. By contrast, the ubiquitination sites themselves (marked with “K” in Fig. 4B) show varying levels of mutational sensitivity, consistent with experimental evidence suggesting their dispensability – ubiquitin does not necessarily need to be added to these lysines for proper regulation [51]. In addition, ProteoCast analysis highlighted four cysteines, namely C710, C731, C786 and C802, in the mutational sensitivity profile (Fig. 4B). These predictions, along with experimental studies supporting the essential regulatory role of cysteines within intrinsically disordered regions in response to Reactive Oxygen Species (ROS) [52, 53], suggest this region may function in a redox-dependent manner, with conditional disorder [54].

### Case study on Toll pathway proteins

We finally conducted a focused analysis to examine the relationship between natural polymorphism, available cross-species sequence data, and ProteoCast predictions, using a multi-protein case study from the Toll pathway (Fig. 5, Table S6). This pathway is essential to the developmental patterning of the dorso-ventral axis as well as immunity. Consistent with our proteome-wide analysis, ProteoCast missclassified only a small fraction of population polymorphisms (48 out of 344) in these proteins. The Toll membrane receptor, for example, exhibits 54 mutations in the DEST2 and DGRP populations, all but 4 deemed neutral by ProteoCast. The vast majority of the misclassified mutations (38 out of 48) are exclusively observed in the DEST2 pool (Table S6). None of the mutations exclusively observed in the DGRP are predicted as impactful. This result highlights the purifying selection operating on flies inbred in the laboratory for more than 20 generations - highly homozygote - compared to wild individuals - mostly heterozygote. Moreover, mutations observed exclusively in the DEST2 pool are about four times more numerous than those exclusively observed in the DGRP, in line with proteome-wide estimates (DEST2/DRGP ratio of 3.96 for all 22,392 representative proteoforms). Larger proteins tend to harbour more polymorphisms, due to more positions being mutated while the per-residue polymorphism rate remains constant (mean = 1.04, s.d. = 0.06), a tendency also confirmed at the level of the entire proteome (adjusted R^2^ = 0.33 for 18,238 proteoforms spanning 40-1,000 residues).

**Figure 5.**
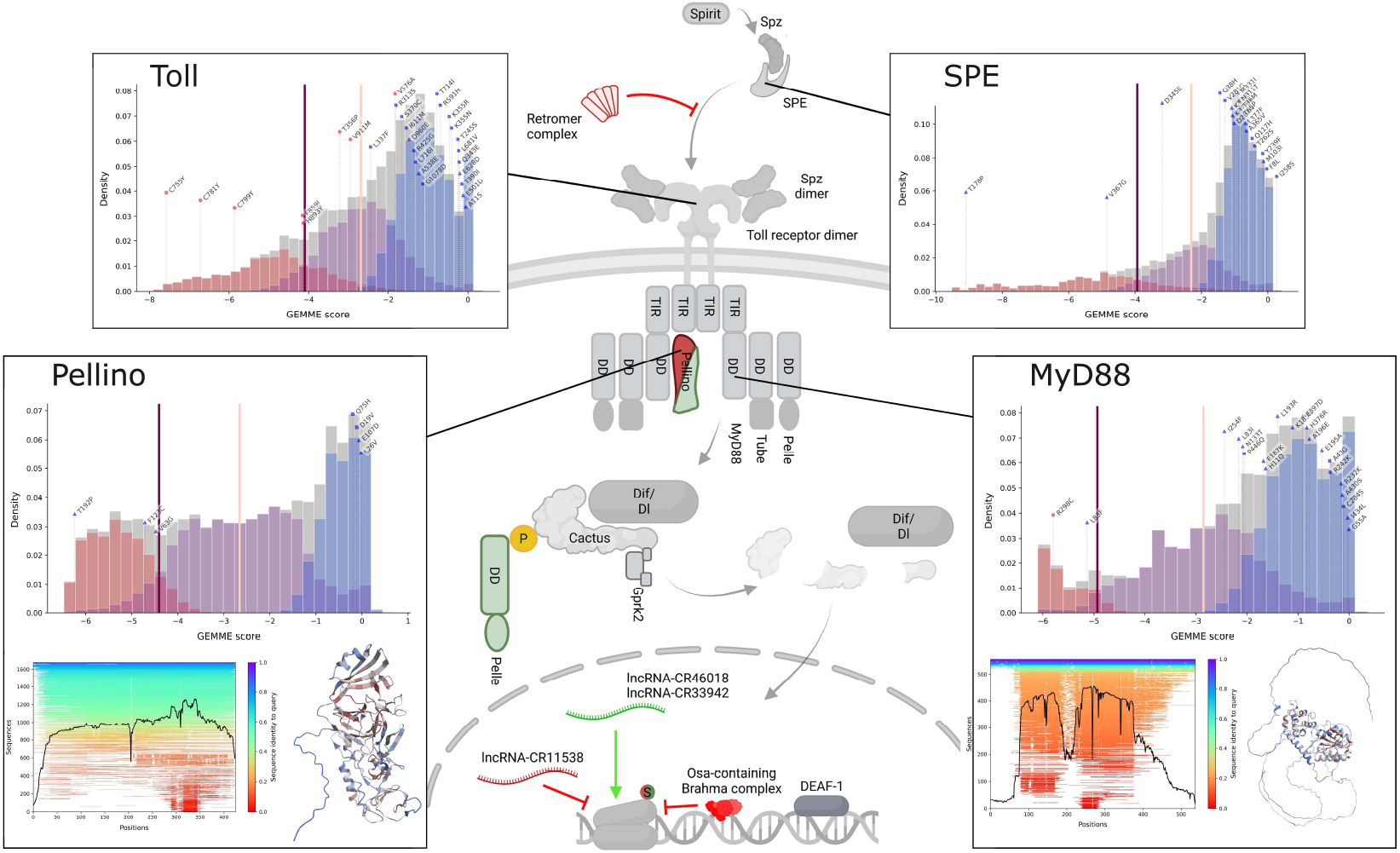
ProteoCast predicted mutational landscape for proteins from the Toll pathway. ProteoCast score distributions and variant classification are represented for four proteins of the Toll pathway, the membrane receptor Toll (FBpp0084431, 1,097 residues), the serine protease SPE (FBpp0083832, 400 residues), the transducer MyD88 (FBpp0087679, 537 residues) and its negative regulator Pellino (FBpp0083913, 424 residues). The three fitted Gaussians are highlighted in colors and the vertical bars indicate the variant classification thresholds, as in Fig. 2B. Sampled natural polymorphisms (inbred or wild) and reference lethal mutations are depicted as blue and red points, respectively. For Pellino and MyD88, we also show the input alignment plots and the AlphaFold-predicted 3D models colored according to ProteoCast-predicted mutational sensitivity, from blue through white to red. The schematic representation of the Toll pathway is adapted from Valanne et *al*. [55].

The fraction of mutations observed in the input alignment varies widely across proteins, from less than 10% for Mtk – suggesting its specificity to a few organisms – to over 90% for the extracellular serine proteases Snake and SPE, both cleaving the Toll ligand Spaetzle. This high fraction reflects the ubiquity of this family of enzymes in eukaryotes and prokaryotes. Intermediate fractions (50–65%) were observed for both the intra-cellular proteins MyD88, a transducer of the Toll receptor, and its negative regulator Pellino. However, the mutational landscapes of these two proteins contrast strikingly. MyD88 contains unstructured regions that are poorly covered in the input alignment and prone to accumulating natural polymorphisms (19 out of 21 classified as neutral), and thus less likely to alter the protein function. Conversely, Pellino is largely structured and the alignment is dominated by homologs sharing high sequence similarity (*>*40%). Its low number of natural variants in DEST2 (5 mutations) likely stems from strong structural constraints.

Overall, the systematic application of ProteoCast can assist experimentalists and theorists in exploring the role of natural polymorphism in the function of specific proteins or entire pathways, offering complementary information for interpreting Genome Wide Association Studies (GWAS).

### Comparison with other state-of-the-art methods

Having demonstrated ProteoCast’s accuracy on Drosophila-specific benchmarks and case studies, we further assessed its broader applicability through independent evaluation on standard benchmarks (Table 2). We evaluated ProteoCast without any parameter adjustment against established methods originally developed using these or similar datasets.

**Table 2.** Independent evaluation of ProteoCast on standard benchmarks compared to established methods. For each dataset, we indicate the ProteoCast functionality evaluated, the number of data points, the comparison methods used, whether these methods are supervised, and whether the evaluation benchmark was used in the original method publications (indicating potential use during development). Main results are summarised with references to corresponding tables and figures. *the methods were trained on other CAID versions or other binding datasets.

| ProteoCast functionality | Evaluation benchmark | Comparison methods | Supervised (Y/N) | Used the benchmark (Y/N) | Results summary |
| --- | --- | --- | --- | --- | --- |
| Variant classification | ClinVar [56] (63k) | EVE [8] | N | Y | higher coverage for ProteoCast, similar AUC-ROC for both methods, slightly higher recall for EVE on a variant subset (Table S7, Fig. S15) |
| Functional site identification in unstructured regions | curated set from the yeast proteome [57] (526) | phylo-HMM [57] | N | Y | substantially higher detection rate for ProteoCast (Table S9, Fig. S16) |
|  | CAID3 Binding-IDR [58] (64) | top-10 CAID3 predictors | Y | N* | ProteoCast on par with the other predictors (Tables S10 and S11, Fig. S18) |

#### Distinguishing pathogenic from benign variants in ClinVar

ProteoCast provided confident predictions for 99% of the 63,000 human variants annotated in ClinVar, correctly identifying 77.1% of the pathogenic ones as mild or impactful with a specificity of 86.9% (Table S7). Moreover, it prioritised some pathogenic variants among the top 10 strongest predicted effects or among the top 5 most sensitive residues for 632 out of 2,525 proteins. For instance, ProteoCast identified the disease mutation R578H, associated with Pitt-Hopkins syndrome, as the most impactful variant in transcription factor 4 (TCF4) among 12,749 possible substitutions. These results demonstrate that ProteoCast effectively captures human genetic and disease patterns. For comparison with the state of the art, we chose EVE, which, like ProteCast, performs unsupervised variant classification based on predicted evolutionary raw scores [8]. We made the comparison on the subset of 1,736 human proteins from ClinVar that are covered by EVE (data downloaded from https://evemodel.org/download/bulk). EVE provided predictions for only 88% of the 47,022 variants annotated in these proteins based on its input MSA quality assessment. This assessment is conceptually similar to our confidence score but results in a much lower coverage of the mutational landscape for EVE (median of 80%, minimum of 5%) compared to ProteoCast (median of 99% and minimum of 79%, Fig. S15B). EVE further excluded 25% of the remaining variants because of high uncertainty on its predictions (Fig. S15C-D). This protocol led to lower recall (69.0%) and specificity (72.5%) compared to ProteoCast (78.3% and 85.2%) on the full dataset (Table S7). Both EVE and ProteoCast achieved similar specificity (*∼*88) on the subset of 33,274 variants confidently classified as pathogenic or benign by EVE, with a slightly higher recall for EVE (86.6 versus 81.9 for ProteoCast, Table S7). Overall, ProteoCast provides a good balance between sensitivity and specificity across virtually all mutations, while EVE, despite achieving higher recall on the subset it classifies, leaves a substantial fraction (20%) of annotated variants uncertain.

#### Identifying functional sites in the disordered fraction of the yeast proteome

We focused on the curated set of 500 phosphorylation sites, localization signals, degradation signals, SUMO sites, and interaction motifs defined in Nguyen Ba et *al*. (2012) (see Table S8). Half of the sites are shorter than 4 residues, with a maximum length of 65 residues (Fig. S17A). Among the 437 low-pLDDT sites, ProteoCast fully included 186 sites within segments showing locally elevated mutational sensitivity —more than twice the number detected by phylo-HMM (99 sites, Table S9). When considering the 347 sites labeled as disordered by [57], ProteoCast achieved detection rates of 38% compared to 30% for phylo-HMM (Table S9). These results demonstrate that modeling mutational sensitivity as a continuous signal from large sequence datasets allows recovering evolutionary constraints that cannot be detected by phylogenetic HMMs over a dozen closely related species. For instance, in SCD5, phylo-HMM detects only one out of three copies of the motif “LKPTATG”. Despite the relatively low conservation of the two others, ProteoCast contrasts them with high precision against their background (Fig. S16).

#### Identifying disordered binding sites from CAID3

The 62 annotated binding sites in CAID3 [58] dataset (https://caid.idpcentral.org/assets/sections/challenge/static/references/3) are substantially longer than PTM sites or SLIMs, with a median length of 19 residues and extending up to 989 (Fig. S17C). 40% of them were fully included in ProteoCast segments with elevated sensitivity (Table S10). Using sensitivity differences between the identified segments as a predictive metric, we achieved residue-wise performance comparable with the top 10 performing predictors in the CAID3 challenge – our approach is ranked 6th based on AUC (Fig. S18 and Table S11). Relying solely on per-residue mutational sensitivity yielded lower performance. These results demonstrate that our unsupervised MSA-based approach identifies disordered binding sites as effectively as supervised machine learning methods trained specifically on this type of data, including those leveraging embeddings computed by pre-trained protein language models.

## Discussion

### Biological insights from Drosophila proteome analysis

Proteocast has three primary purposes for the Drosophila community and beyond. Firstly, it facilitates the identification of genome-editing targets by enabling rapid prioritisation of the hundreds of millions of possible missense mutations over the fly proteome. Secondly, it sheds light on the evolutionary conservation and 3D structural context of hundreds of thousands of missense variants found in inbred and wild fly populations and lacking annotations. Thirdly, it can aid in elucidating the role of unstructured regions of the fly proteome by pointing at potential interaction and regulatory sites within these regions. It offers freely accessible, user-friendly and interactive online web services for a detailed examination of any single-point variant of interest at https://proteocast.ijm.fr/drosophiladb/.

We have provided functional validations of our prediction using the rich datasets available for Drosophila as well as CRISPR experiments. These results demonstrated ProteoCast’s ability to accurately predict organism-level mutational impact, specifically in terms of binary phenotypic outcomes such as developmental viability. Moreover, they allowed for discovering three recessive developmentally lethal mutations in a protein involved in the biosynthesis of a coenzyme central to metabolism. Ongoing and future work will aim at full biochemical characterisation of the generated mutants.

Furthermore, our analysis has revealed that a substantial fraction of variants found in inbred and wild fly populations are counter-selected in evolution, suggesting that they may alter protein or organismal fitness. Some of them occur in proteins with known lethal missense mutations, thus likely essential for the organism. For instance, ProteoCast identified impactful DEST2 mutations clustered near the putative DNA-binding interface of the nucleosome remodeler Mi-2, and within the kinase domain of the Doa somatic sex determination protein, where the natural variant V364G scored more severely than the nearby lethal mutation C275Y (Fig. S19). These findings call for biochemical characterisation of the putative impact of these naturally occurring mutations on protein activity and interactions. More broadly, these observations raise questions about how such potentially deleterious variants are maintained differently across population types. ProteoCast predictions could facilitate multiple comparative approaches: examining the differences between the polymorphism fixed in DGRP, inbred in the laboratory for more than 20 generations (F = 0.986), with that of natural populations in DEST2, or studying seasonality, geography and natural polymorphism by focusing on impactful DEST2 variants rather than genome-wide scans. We explored the latter approach for two xenobiotics detoxifying genes, Cyp6a9 and Cyp317a1, with geographically influenced incidence (Fig. S20). Reciprocally, while ProteoCast helps interpret natural variants, these population data may in turn help refine annotations for the fly proteome. We identified 32 mutations referenced as developmentally lethal in FlyBase, yet observed in inbred or wild fly populations. ProteoCast provided high-confidence predictions for 21 mutations, classifying all but one as neutral, highlighting the need for better curation of lethality annotations. For example, the R10K mutation in hopscotch kinase is annotated as lethal but likely owes its phenotype to an accompanying premature stop codon that eliminates the entire kinase domain [59].

Overall, the evolutionary data available for the fly allowed obtaining high-resolution for most of the proteins. A subset of 725 unique proteoforms from 593 genes even have their mutational landscapes nearly entirely covered by the input alignment, with more than 95% of all possible substitutions actually observed in natural sequences (Fig. S21). This natural mutational saturation highlights the ubiquity of the corresponding protein families across life kingdoms. These proteoforms have many identifiable homologs (at least 4,000 sequences in the alignment) that retain detectable similarity while exhibiting extensive variation at individual sites. They are more likely to harbour mutations impairing organismal fitness compared to the rest of the proteome - 1.8 fold enrichments in lethal mutations. They tend to be more connected in the fly interactome, with a 2.8-fold higher likelihood of having *>*20 protein partners. Furthermore, their genomic loci are significantly enriched for regions associated with transcription factor (TF) binding sites recognised by early development genes such as *CTCF, BEAF-32* or *Dref* that are also involved in chromatin domain organisation and early development (**Fig. S22**). Collectively, these findings suggest the functional importance of these proteins to the organism through their involvement in key developmental processes, and that such importance can, to some extent, be inferred directly from its evolutionary history, without needing additional information about population, genetics or interactomics data.

Turning to structural applications, we systematically mapped ProteoCast predictions on AlphaFold-predicted 3D models over the fly proteome. While both methods exploit multiple sequence alignments, ProteoCast extracts evolutionary signals unsupervised whereas AlphaFold2 maps alignments to 3D coordinates. Regions with low pLDDT (indicating mapping difficulties) can still harbour sites under selection detectable by ProteoCast, thus remaining important for protein interactions and functions. Our results demonstrate ProteoCast’s potential for transparent screening of phosphoproteomics datasets to identify functionally relevant PTM sites without expert intervention. Its adaptive detection of evolutionary signals in unstructured regions through an automated segmentation may prove advantageous over methods using absolute thresholds [60]. Beyond filtering known PTM sites and linear motifs, ProteoCast can discover uncharacterised functional sites in disordered regions.

### Methodological innovations and comparative advantages

This study builds upon established approaches for accurate, interpretable, and scalable prediction of variant effects that exploit evolutionary information from protein sequences [7, 31]. ProteoCast introduces several methodological innovations compared to GEMME by (i) integrat-ing evolutionary scores with protein structural features, (ii) identifying regulatory sites within unstructured protein regions, and (iii) providing calibrated confidence metrics and systematic variant classifications to guide experimental prioritisation. While individual methods in the field address some of these aspects separately, ProteoCast uniquely integrates all three within a single evolutionary-based framework.

ProteoCast provides a graduated classification of variants as neutral, mild, and impactful, aimed at helping users interpret the predicted mutational landscape and prioritise variants for experimental validation. This classification encompasses all possible variants of a given protein, in marked contrast with approaches like EVE that use GMM probabilities to estimate uncertainty and exclude uncertain variants from evaluation. Our approach is pragmatic and transparent, avoiding potential model or dataset biases that may lead to inconsistencies between statistical probabilities and actual biological uncertainties. ProteoCast further leverages its continuous mutational sensitivity scores through a segmentation strategy designed to identify functional motifs including regulatory and binding sites in unstructured protein regions. This strategy represents a substantial conceptual advance over previous foundational works for addressing this problem [57, 61].

First, we exploit mutational sensitivity rather than conservation, as we observed that the former provides higher resolution and better agreement with experimental measurements of protein stability [30]. Second, our segmentation approach models mutational sensitivity as a piecewise constant continuous parameter, providing a quantitative framework for capturing and disentangling different levels of evolutionary constraints across protein segments. As Figure 4 clearly illustrates, mutational sensitivity contains substantially richer information than binary signals, and our model is considerably more expressive than conservation-based models relying on binary state distinctions [57, 61]. This conceptual framework is supported by previous studies demonstrating the superior accuracy of segmentation approaches over simpler “two-state” models on diverse biological datasets, including RNA-seq [62] and ChIP-seq data [63], reinforcing their suitability for capturing the complexity and heterogeneity of biological signals.

### Limitations and perspectives

Despite the robustness of our approach, alignment errors might result in variant or residue misclassification. Protein extremities are typically highly gapped in the input alignment and ambiguities in gap opening/closing positions may arise at the borders of poorly covered regions. Nevertheless, ProteoCast confidence metrics provide a means for flagging potentially problematic positions. For instance, half of the 22 lethal or hypomorph mutations affecting the first methionine are misclassified as neutral, but most of these incorrect predictions (7/11) are flagged as unreliable due to poor alignment quality. This demonstrates that our confidence metrics effectively identify problematic predictions. More broadly, while alignment quality parameters and thresholds might theoretically be optimized differently for distinct applications, our approach prioritises practical usability with minimal parameter tuning. Our validation across diverse tasks and the entire Drosophila proteome demonstrates the robustness of using standardized alignment generation protocols for evolutionary constraint-based analyses. Future versions of ProteoCast will explore integrating complementary approaches based on protein language models (pLMs) to further improve prediction accuracy in challenging regions. This is a promising avenue as we previously showed that the pLM-based predictor VespaG helps to regularise GEMME scores thanks to its knowledge of a universal protein representation space [12].

Future research will aim at a tighter integration of evolutionary and structural data. For instance, applying ProteoCast segmentation strategy to AlphaFold2 pLDDT profiles revealed that around 34% of the known PTM sites and 35% of the known SLiMs fall within regions modeled with higher confidence than their surroundings. This detection likely reflects AlphaFold2’s ability to recapitulate and generalise experimental knowledge about conditional folding. It only partially overlaps with ProteoCast detection from the predicted mutational sensitivity profiles, suggesting avenues for combining the two types of information. Another direction will consist in extending the resource to other common model organisms, such as *X. laevi, C. elegans, S. cerevisiae*, and *E. coli*, as well as *H. sapiens*. The wealth of mutational data and associated phenotype will help further assess the quality of our predictions. The generated data will facilitate the evaluation of large-scale predictions against phenotypic knowledge at both organismal and molecular levels as well as significantly ease the identification of residue-specific genome editing. Our findings will serve as a starting point toward identifying the molecular determinants of continuous phenotypic traits controlled by multiple genes.

## Methods

### Drosophila Experiments

#### Lines used

*da*GS driver [64] (provided by the Monnier lab, Université de Paris Cité).

#### Vienna Drosophila Resource Center stock (gene name)

42571 (*JIL-1*, FBgn0020412), 31712 (*brm*, FBgn0000212), 31053 (*HDAC6*, FBgn0026428), 61875 (*Naprt*, FBgn0031589), 35704 (*smid*, FBgn0016983), 67758 (*Gpo3*, FBgn0028848), 31395 (*Rm62*, FBgn0003261), 28038 (*nAChRβ2*, FBgn0004118), and 27503 (*CadN*, FBgn0015609).

#### RNAi-induced developmental lethality

All the flies are kept in closed vials in incubators at controlled temperature, humidity (60%) and 12 hours light cycle. Experiments are carried at 26^*°*^C. 5 individual replicate vials were made per condition. Each RNAi was induced using two distinct concentrations of RU486 0 *µ*g/mL (negative control) and 20 *µ*g/mL, the maximum inducer concentration usable during development as previously described [65]. Each contains 6 *da*GS virgin females and 5 RNAi males left to cross and lay eggs for 48 hours on the following food composition: 8.33% (w/v) yeast, 8.33% (w/v) corn, 1.13 (w/v) agar and Methyl 4-hydroxybenzoate (Moldex) at a final concentration of 5.3 g/L to prevent fungal contamination. Eclosing flies were then counted daily for 10 days.

### Datasets

#### FlyBase

We retrieved the *Drosophila melanogaster* proteome version 6.44 as a FASTA file (*dmel-all-translation-r6.44.fasta*), available at FlyBase [34] (https://ftp.flybase.net/genomes/). This transcriptome contains a total of 30,738 sequences, of which 22,392 are unique. We focused our attention on single point mutations generated by ethyl methylsulfonate (EMS) and made a list of those associated with developmental lethality - using the Phenotype QuickSearch tool provided by Flybase - as well as a second list of hypomorph mutations. We filtered out the 250 mutations overlapping between the two.

#### Drosophila Genetic Reference Panel

We obtained all polymorphisms from the 205 inbred lines of the Drosophila Genetic Reference Panel (DGRP) [32] in a text file (*dgrp.fb557.annot.txt*). This dataset contains approximately 4.5 million genomic variations, including deletions, insertions, single nucleotide polymorphisms (SNPs), and multi-nucleotide polymorphisms (MNPs). SNPs are the most prevalent type of variations (89% of all detected variants), 4.5% of which result in missense mutations leading to amino acid changes in proteins. Mapping these missense mutations to the fly proteome sequences resulted in 178,264 unique variants affecting 20,137 (89.9%) proteoforms from 12,682 genes (90.8%).

#### Drosophila Evolution over Space and Time

The Drosophila Evolution over Space and Time (DEST) [33] resource is a large-scale dataset that compiles genomic polymorphism data from natural *Drosophila melanogaster* populations sampled across diverse geographic locations and time periods. We obtained all annotated SNPs from the August 2024 version of the dataset in VCF format (*dest.all.PoolSNP.001.50.3May2024.ann.vcf.gz*). It includes approximately 4.8 million SNPs classified into 27 different annotation types. For this study, we focused specifically on missense SNPs, identifying 332,848 variants that affect 94% of Drosophila genes and 92.4% of proteoforms. After resolving mismatches in gene and transcript identifiers, we retained 285,771 unique missense SNPs for further analysis.

#### Post-translational Modifications

We retrieved all PTM sites annotated for the Drosophila melanogaster proteome in the iProteinDB [43] database at https://www.flyrnai.org/tools/iproteindb/through its API. We identified PTM sites in 8,530 representative proteoforms (38%) from 4,276 genes (31%). The vast majority of these sites are targeted by phosphorylation (*∼*92%), the remaining sites being oxidized or acetylated.

#### Short Linear Motifs

We obtained 113 short linear motifs (SLiMs) ranging from 3 to 20 amino acids, from the Eukaryotic Linear Motif (ELM) resource (http://elm.eu.org/) [48]. These motifs were retrieved from a downloadable file, *elm instances.tsv*, using the following search path: ELM DB *→* ELM instances *→*Filter by organism (*Drosophila melanogaster*).

### ProteoCast workflow

The ProteoCast workflow unfolds in five main steps.

#### 1. Multiple sequence alignment generation (Fig. 1A)

Given an input query protein sequence, it retrieves related sequences using the MMseqs2-based protocol implemented in ColabFold [37].

#### 2. Full single-mutational landscape prediction (Fig. 1A)

It uses the variant effect predictor GEMME [7] to estimate the impact of all possible amino acid substitutions at every position in the query.

#### 3. Prediction reliability assessment (Fig. 1B)

It flags the query as having unreliable predictions due to insufficient evolutionary information if the input alignment contains less than 200 sequences. Otherwise, it assigns binary confidence scores that reflect alignment quality and diversity to each residue in the query.

#### 4. Variant and residue classification (Fig. 1B)

It classifies variants as impactful, mild or neutral by fitting a Gaussian mixture model [66] to the predicted raw scores. It further classifies residues as sensitive to mutations if at least 10 out of the 19 possible substitutions are mild or impactful. Otherwise, residues are considered tolerant to mutations.

#### 5. 3D structural mapping and segmentation (Fig. 1B)

It maps residue mutational sensitivity scores onto the 3D model predicted by AlphaFold2 for the query protein. It decomposes the mutational sensitivity profile into segments using the FPOP algorithm [46, 47]. The granularity of the segmentation accounts for AlphaFold pLDDT profile.

In the following, we give details about the different steps, illustrating their application over the entire fly proteome.

### Step 1. Multiple sequence alignment generation

For every unique proteoform from the fly proteome (FlyBase version 6.44), we generated a multiple sequence alignment (MSA) using the ColabFold (CF) protocol [37]. We previously showed that this protocol compares favourably with workflows relying on profile hidden Markov models for the prediction of mutational effects [31]. It is much faster, thanks to its usage of the highly efficient sequence MMseqs2 search algorithm [36], and it yields similar or better predictive accuracy. CF protocol produced two alignments for each input sequence by searching against UniRef30, a clustered version at 30% sequence identity of UniRef100 [67], and a curated database comprising various environmental sequence sets [37] (ColabFold environmental database, ColabFold Env). We used UniRef30 v2103 and ColabFold Env v202108. We concatenated the alignments and gave the resulting one as input for GEMME. The CF protocol did not manage to find any homolog for 9 proteoforms.

### Step 2. Full single-mutational landscape prediction

We utilised the *reformat.pl* tool from the HH-suite [68] to convert A3M alignment files into FASTA format. We handled sequences containing undefined residues (represented by symbols “X” or “U”) by employing the following strategy: when the undefined residue was located at the beginning of the sequence, the rest of the corresponding column in the alignment was always filled with gaps, and thus we removed that column. For other instances of undefined residues, we replaced them with the most frequently occurring amino acid at the respective position(s) within the alignment. We ran GEMME with default parameters except that (1) we increased the memory (RAM) to handle very long proteins (*Xmx9126m*), (2) we increased the number of JET [69] iterations to 7 to ensure robustness. The JET method subsamples the initial input set of sequences, generates a tree for each subset and determines each position’s evolutionary trace, reflecting the level in the tree where an amino acid appeared at that position and remained fixed thereafter. It estimate a conservation value *T*_JET_(*s*^*∗*^, *i*) for each residue *i* in the query *s*^*∗*^ by averaging evolutionary traces over the generated trees. Subsampling is performed using a Gibbs sampling strategy and hence, multiple runs may produce different conservation estimates. We found that the Pearson correlation between the conservation values computed from two independent runs could be as low as 0.78 over a subset of 200 protein sequences. Setting the number of iterations to 7 and retaining the maximum conservation value for each residue improved reproducibility, with correlation coefficients systematically above 0.95 for any two runs. Another parameter in JET controls the number of sequences from the input alignment that are used to estimate residue evolutionary conservation. By default, it is set to 20,000 sequences, a threshold that effectively allows us to retain all sequences for the vast majority (97%) of proteoforms. In total, GEMME produced some predictions for 22,169 unique proteoforms (Table S1).

### Step 3. Prediction reliability assessment

A popular measure for estimating the diversity of an alignment is the number of effective sequences [9, 25, 70]. It is computed by summing up sequence-specific weights that reflect their overall similarity. However, we previously showed that this metric is not a good indicator of the quality of GEMME predictions [31]. Instead, we found that ColabFold’s MSA generation protocol, which maximises diversity and minimises the number of sequences, allows to set a reliability threshold based on the number of sequences in the MSA. Building on these previous results, ProteoCast flags proteins with fewer than 200 homologous sequences in the input alignment as likely to have unreliable predictions. For the remaining proteins, it estimates a binary-valued confidence score for each residue. Instead of simply computing the percentage of gaps in the alignment, as is often done, ProteoCast metric integrates this measure with the dispersion of predicted scores, the estimated conservation, and the fraction of substitutions actually observed in the alignment. This multifactorial approach better captures the complexity of alignment quality than simple gap counting or sequence diversity metrics alone. The confidence score *c*(*s*^*∗*^, *i*) for residue *i* in the query sequence *s*^*∗*^ is expressed as,

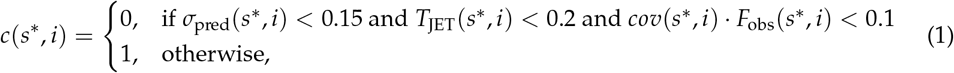

where *σ*_pred_(*s*^*∗*^, *i*) is the standard deviation of the 19-long vector of predicted variant effects, *T*_JET_(*s*^*∗*^, *i*) is the conservation value, *cov*(*s*^*∗*^, *i*) is proportion of sequences that have an amino acid at position *i* (as opposed to a gap), and *F*_obs_(*s*^*∗*^, *i*) the fraction of observed substitutions at position *i* expressed as,

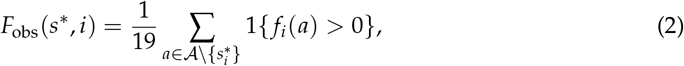

where *f*_*i*_(*a*) is the frequency of occurrence of the amino acid *a* at position *i* in the MSA. The resolution of the predicted mutational landscape directly depends on *F*_obs_(*s*^*∗*^, *i*). Indeed, GEMME computes an evolutionary distance between the query sequence and the closest sequence in the input alignment displaying the mutation of interest. Mutations not observed in the alignment are assigned the maximum evolutionary distance.

#### Threshold tuning

We assessed the influence of changing the thresholds for *σ*_pred_(*s*^*∗*^, *i*), *T*_JET_(*s*^*∗*^, *i*), and *score*(*s*^*∗*^, *i*) on the number of segments *N*_seg_ detected as unreliable (low confidence score) and the proportion of segments of length one (isolated unreliable residues, *f*_seg1_). Wetested several combinations of threshold values in the intervals: *σ*_pred_ *∈{* 0.1, 0.11, 0.12, 0.13, 0.14, 0.15 ; *}T*_JET_ *{* 0.1, 0.15, 0.2, 0.25, 0.3 ; *} score ∈ {* 0.05, 0.1, 0.15, 0.2 *}* (Fig. S23). We obtained very similar results for the tested combinations, indicating that they are robust within these value ranges. We observed a slight tendency for fewer segments and relatively more single-residue detections upon increasing the thresholds (Fig. S23).

#### Smoothing

We further smooth the per-residue confidence scores over the sequence to mitigate the discontinuities that may arise from using deterministic cutoffs. The final confidence score for the *i* ^th^ residue of the query sequence *s*^*∗*^ is thus computed as:

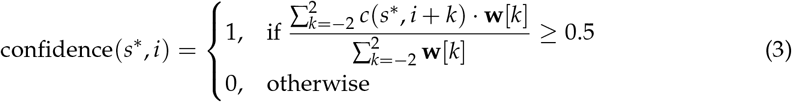

where the weight vector **w** = [2, 3, 4, 3, 2] (Fig. S24).

### Step 4. Variant and residue classification

The range of variant predicted raw scores may vary substantially from one protein to another, making direct comparisons over the entire proteome difficult. This motivated us to devise an adaptive variant classification strategy tailored to each protein-specific raw score distribution. Namely, ProteoCast uses a Gaussian Mixture Model (GMM), a probabilistic approach that assumes the data are generated from a mixture of Gaussian distributions [66]. The GMM parameters are estimated through the Expectation-Maximization (EM) algorithm [71]. Similar strategies with two or three Gaussians were previously proposed to classify experimental measurements [28] or scores predicted by a variational auto-encoder [9].

We used the mixture module from python *scikit-learn* library [72] with component-specific general covariance matrices (“full” type), 30 initialisations, a convergence threshold of 10^*−*5^, and a maximum number of EM iterations of 1,000. We quantified the effect of the number of Gaussian components on model fit across 20 trials of 100 randomly selected proteoforms. We excluded scores with low MSA-based local confidence to reduce risk of bias they might introduce. For each number of components *k* = 1…5, we computed the mean Akaike Information Criterion (AIC) and the log-likelihood (LL) per trial, normalised by the number of data points per protein. AIC balances model fit (*i.e*., the log-likelihood) and model complexity (number of parameters) and thus penalises overfitting. Both metrics indicate improved model quality with increasing components (Fig. S3). Log-likelihood increases by 7% and AIC decreases correspondingly from 1 to 2 components, and by 2% from 2 to 3 components, on average. All proteins benefit from the increased number of components up to three. Beyond three Gaussians, the average gains become negligible (*<*1%) and the goodness-of-fit even deteriorates for a substantial fraction of proteins, and hence, we fixed the number of components to three.

Given the three identified components, G1, G2 and G3, with mean values *µ*_*G*1_, *µ*_*G*2_, and *µ*_*G*3_, respectively, and *µ*_*G*1_ *< µ*_*G*2_ *< µ*_*G*3_, ProteoCast defines three variant classes as follows,

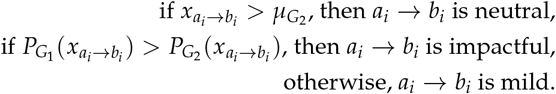

with 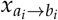 be the raw score predicted for substituting the amino acid *a* by *b* at position *i*. Moreover, for each variant, ProteoCast computes a Z-score relative to the distribution of neutral mutations, Z-score 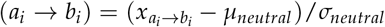, where *µ* and *σ* stand for the mean and standard deviation, respectively.

### Step 5. 3D structural mapping and segmentation

For every unique proteoform from the fly proteome (Flybase version 6.44), we retrieved its 3D model either by batch downloading (https://www.uniprot.org/proteomes/UP000000803) from the AlphaFold Database (13,475 structures), or by using Python *requests* library [73] (5,119 structures). For the missing structures, we generated all those with fewer than 4,000 amino acids (3,345 structures) by running ColabFold with default parameters. ProteoCast then mapped the computed per-residue mutational sensitivity values (average of raw scores) and classes onto each protein’s 3D model. It further used AlphaFold pLDDT scores to segment the mutational sensitivity profile, as described in the following.

The problem of segmenting a one-dimensional continuous measure along a protein sequence can be formulated as follows. We consider *D* changepoints *τ*_1_ *<* … *< τ*_*D*_ within the range 1 and *n −* 1. These changepoints correspond to unknown positions along a protein of length *n*, where a shift in the mean of the measure is observed. We adopt the convention that *τ*_0_ = 0 and *τ*_|*τ*|_ = *n*. These changepoints define |*τ*| = *D* + 1 distinct segments. The *j*-th segment includes the data *{τ*_*j−*1_ + 1, …, *τ*_*j*_ *}*. Each segment is premised on the assumption that the *Y*_*i*_ therein are independent and follow the same Gaussian distribution, with a mean *µ*_*j*_ specific to that segment and a common variance *σ*^2^. Expressed mathematically, we have:

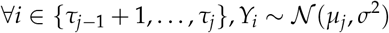

The parameters of the model, including *τ*_1_ *<* … *< τ*_*D*_, can be estimated using penalized maximum likelihood inference. To achieve this, we used the FPOP algorithm [46, 47] which solves the inference problem exactly with dynamic programming. Informally, the FPOP algorithm works by considering the penalized maximum likelihood of the data from observation 1 to *t* as a function of the parameter (the mean) of the last segment. This concept is referred to as ‘functional pruning’. In the Gaussian case, the resulting function is piecewise quadratic. When a new observation at time *t* + 1 is made, this function can be efficiently updated (i.e., compute the penalized likelihood penalized maximum likelihood function from observation 1 to *t*) using a formula similar to that of the Viterbi algorithm. This formula is applied piece by piece or interval by interval. At each step, the algorithm searches for the optimal value of the parameter for the last segment that maximizes the penalized likelihood.

The number of changepoints *D* estimated by FPOP is a decreasing function of the penalty *ασ*^2^ log(*n*). The constant *α* is a hyperparameter set to 1.4, determined after assessing the fraction of detected PTMs in the resulting segmentations while ensuring the visual goodness-of-fit (Fig. S25). The variance *σ*^2^ is estimated on the data using the Hall estimator [74]. We observed that the variations in mutational sensitivity were higher in protein regions modelled with high confidence by AlphaFold2 (pLDDT *>* 70) compared to other regions. To avoid over-segmenting these areas (by introducing artificial changepoints), we applied lower weights to the observations in these regions, with *w* set to 0.1, while keeping it at 1 elsewhere. After estimating the segments, we assign discrete scores to each of them. A score of 0 indicates that the means of both adjacent segments are greater than that of the current segment; a score of 1 that one adjacent segment has a lower mean and the other a higher mean than the current segment; and a score of 2 that the means of both adjacent segments are lower than that of the current segment.

### Implementation and computational details

To generate the input we run ColabFold protocol, multiple sequence alignment generation part, through the following command: `colabfold_search --threads=8 --db-load-mode=2 $fasta_file $ {COLABFOLD-DB} $out folder`. For a set of 3,000 sequences comprising a total of 1,185,789 amino acids it took 1.5 hours to generate MSAs on the supercomputer “*MeSU*” of Sorbonne University (16 CPUs from AMD EPYC Milan processors, 400GB shared RAM memory).

We implemented ProteoCast in Python and R. ProteoCast relies on *scikit-learn* library [72] to fit a Gaussian mixture model, the *gemmi* library [75] to parse PDB files and manipulate the B-factors associated to the structure, the *re* module to parse annotation data files, and *pandas* for data manipulation. For the same set of the protein sequences the generation of mutational landscapes, classification and analysis (Fig. 1A *Mutational Landscape* and Fig. 1B) took 4 hours on 40 CPUs.

The online database (https://proteocast.ijm.fr/drosophiladb/) is a *Django*-based project running on a virtual machine hosted by *Institut Jacques Monod* (Dell PowerEdge R740 server). The backend was developed in Python using *Django*, while the frontend was built with HTML, CSS, and JavaScript to provide a user-friendly interface. The application was deployed on an Apache2 web server with Django’s WSGI integration. It incorporates *plotly* for interactive data visualization and *Mol* (Molstar)*[76] for 3D structure visualisation. Data handling and processing were facilitated using *pandas*.

### Evaluation

We estimated the performance with the balanced accuracy metric, suited to unbalanced datasets and is defined as Balanced Accuracy 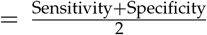 where Specificity 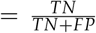in our study the fraction of DGRP or DEST2 polymorphisms predicted as neutral, and Sensitivity (Recall) 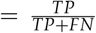in our study the fraction of lethality inducing mutations predicted as impactful. If several proteoforms are affected by a SNP, we take the one with the highest *F*_*obs*_ score.

### Comparison with other methods

#### ClinVar dataset: variant classification and comparison with EVE

We retrieved the ClinVar annotations from the ProteinGym benchmark [25]. They encompass approximately 63,000 missense mutations across 2,525 proteins, each labeled as either pathogenic or benign. For this analysis, we directly took the MSAs and raw GEMME scores we computed in a previous study [31] (https://doi.org/10.5061/dryad.vdncjsz1s) and applied the subsequent ProteoCast analyses to them. For 316 ClinVar sequences that did not match our existing dataset, we computed predictions from scratch with ProteoCast Docker image. As for EVE, we downloaded their bulk predictions from: https://evemodel.org/download/bulk.

ProteinGym provides RefSeq IDs (e.g., NP 001750.1) and sequences for the ClinVar proteins. We performed ID mapping to UniProt accessions for ProteoCast (e.g., P30279) and to UniProt entry names for EVE (e.g., CCND2 HUMAN), using the UniProt ID-mapping tool. EVE predictions were unavailable for 789 proteins due to ID-mapping issues (7), missing predictions for the mapped IDs (506), or sequence mismatches (276), leaving a subset of 1,736 proteins covering 47,022 mutations. Additionally, EVE does not provide predictions for certain residues lacking its internal confidence score, reducing the set to 41,329 mutations (88% of the mapped mutations). If we finally remove the non-confident predictions from ProteoCast, we obtain a final common evaluation set of 41,282 mutations (99.9%).

#### Functional sites in the disordered fraction of *Saccharomyces cerevisiae*

We compared ProteoCast with phylo-HMM on a dataset of 526 yeast protein sites curated from the literature and described in Table S8. These sites were classified into five functional categories: phosphorylation sites (the majority group, representing 82% of all sites), localization signals, degradation signals, SUMO sites, and interaction motifs. We retrieved the 174 corresponding full-length protein sequences thanks to their ORF identifiers (e.g., YKL112W) from the Saccharomyces Genome Database (SGD) archive at http://sgd-archive.yeastgenome.org/sequence/S288C_reference/orf_protein/orf_trans_all.fasta. We slightly adjusted the coordinates provided by Nguyen Ba et *al*. (2012) for five sites, verifying them against the cited literature sources, to obtain exact matches between Table S8 and the SGD FASTA file for all sites. We then ran ProteoCast on all proteins, generating MSAs with the ColabFold protocol (uniref30 2302, colabfold envdb 202108) and using 3D models from AFDB. ProteoCast did not provide predictions for two proteins due to highly gapped MSAs. We subsequently focused on three subsets: (1) 437 sites from 146 proteins displaying average pLDDT*<*70, (2) 347 sites from 113 proteins labelled as “disordered” in Nguyen Ba et *al*., and (3) 333 sites from 112 proteins intersecting sets (1) and (2) (see Table S9).

#### Disordered binding sites from CAID3

We retrieved the CAID3 Binding-IDR dataset (15/07/2025 update) from https://caid.idpcentral.org/assets/sections/challenge/static/references/3/binding_idr.fasta. It comprises 64 annotated binding sites within 52 protein sequences. We excluded two protein sequences, one because it was too short (*<*20 residues) and the other one because the MSA was too gapped. We generated 3D models for all sequences using AlphaFold2 within ColabFold.

CAID3 annotations label each residue as structured (“–”), unstructured (“0”), or unstructured and involved in binding (“1”). Following the official CAID3 protocol, residues annotated as “1” were treated as positives (binding), residues annotated as “0” as negatives (non-binding), while structured residues (“–”) were excluded from the evaluation for a more balanced benchmark and a fairer test of predictor performance.

Binding site identification was then assessed at two complementary levels: segment detection and per-residue prediction. For the first one, segments annotated as states 1 or 2 were considered detected sites (Table S10). For per-residue prediction, designed to enable comparison with other methods providing continuous scores, we used three residue-level metrics: raw mutational sensitivity (ProteoCast-mu residue); segment sensitivity contrast, defined as the maximum difference in mutational sensitivity between a given segment (states 1 or 2) and its two neighbors, set to 0 otherwise (ProteoCast-delta-mu segment); and segment labels mapped to per-residue positives (Table S11 and Fig. S18). Continuous predictions were evaluated using ROC curves, with area under the curve (AUC), average precision (APS), maximum F1 score (F1 max), and balanced accuracy (bac). Metrics were computed with the vectorized cls metrics library (available at https://github.com/marnec/vectorized_cls_metrics.git), as recommended by CAID3.

### Analysis of case studies

We used iCisTarget to search for enrichment in transcription factor (TF) binding sites [77, 78]. In brief, i-CisTarget searches existing databases of known TF binding sites as well as experimental chromatin immunoprecipitation experiments. We considered two categories of genes defined based on the fraction of observed mutations, either *F*_*obs*_ *≥* 95 or 85 *< F*_*obs*_ *<* 90. We extracted a subset of 635 genes for the 85 *< F*_*obs*_ *<* 90 category to have roughly the same cardinality as the *F*_*obs*_ *≥* 95 set (593). We considered the enrichment to be significant with a number of enriched features above 7 for a threshold set at Normalised Enrichment Score (NES) threshold of 5.

We visualised 3D structures with Pymol [79] and we used the *align* command to superimpose structures resolved for homologs. For the Naprt example, after careful examination of the publication accompanying the PDB entry 4YUB [80], we realised that the dimeric arrangement annotated as *Biological Assembly* did not match the one described as biologically relevant in the publication. Hence, we generate symmetry mates with Pymol to identify and visualise the correct arrangement (Fig. S10).

### Database description

For each proteoform, we provide its sequence and the corresponding multiple sequence alignment (downloadable in FASTA format), along with a visual representation, inspired by ColabFold [37]. The mutational landscape can be interactively browsed. It includes raw predicted scores, variant predicted classification, known annotations, and local confidence scores (downloadable in CSV format). Additionally, we provide a visualization of the raw score distribution with variant classification thresholds, and the lethal, inbred, and natural missense mutations mapped on it (downloadable as a CSV file). The segmented mutational sensitivity profile can be interactively zoomed in and out and is accompanied with a color bar for pLDDT values (if a structure is available). Finally, we provide access to the mapping of AF2 pLDDT scores, ProteoCast sensitivity scores and residue classification on the 3D structure (if available, all data downloadable as PDB files with values on the B-factor column).

## Supporting information

all suplementary figures

## Data Availability

The ProteoCast predictions and processed datasets generated in this study have been deposited in the Zenodo repository under accession code 10.5281/zenodo.18488840 [81] [https://zenodo.org/records/18488840]. Interactive visualization and query of the predictions are provided through the ProteoCast web server at https://proteocast.ijm.fr/drosophiladb/. Source data are provided with this paper.

The public datasets used in this study are available from the following repositories: pre-computed gene annotations from FlyBase (https://ftp.flybase.net/releases/FB2022_01/precomputed_files/genes/), variant annotations from DGRP2 based on FB5.57 (http://dgrp2.gnets.ncsu.edu/data/website/dgrp.fb557.annot.txt), variant annotations from DEST2.0 (https://dest.bio/data-files/SNP-tables/dest.all.PoolSNP.001.50.24Aug2024.ann.vcf.gz), and the *Drosophila melanogaster* proteome from AlphaFold DB (https://ftp.ebi.ac.uk/pub/databases/alphafold/latest/UP000000803_7227_DROME_v4.tar). AlphaFold 3D models missing from the proteome archive were retrieved manually from the AlphaFold DB (https://alphafold.ebi.ac.uk/).

## Code Availability

ProteoCast is an open-source software released under the GNU General Public License. The software can be run locally using the Docker image available at https://hub.docker.com/r/marinaabakarova/proteocast/ or accessed through the ProteoCast web server at https://proteocast.ijm.fr/ [82]. All scripts used for data processing, analysis, and figure generation are publicly available on GitHub at https://github.com/abakarovaMarina/ProteoCast [83].

## Acknowledgements

This work was granted access to the HPC resources of the SACADO MeSU platform at Sorbonne Université. Further, thanks to all contributing to open-source programming libraries. Last, not least, thanks to all who deposit experimental data in public databases, to those who maintain these databases, and those who make methods available enriching experimental data. This work has been funded by ADAGIO – Ageing and Natural DeAth GenetIc cOntrollers (ANR- 20-CE44-0010) and co-funded by the European Union (ERC, PROMISE, 101087830). Views and opinions expressed are however those of the author(s) only and do not necessarily reflect those of the European Union or the European Research Council. Neither the European Union nor the granting authority can be held responsible for them. We thank Joël Marchand for providing support with the online database and web platform.

## Author Contributions

EL and MR conceptualised the project. MA, EL, and MR designed the computational framework. MA implemented the ProteoCast framework, performed all analyses and data curation. AL implemented the segmentation algorithm. MA, EL, MR, and AL analysed the results. MA and MIF developed the web platform. MA, EL, and MR wrote the manuscript. All authors reviewed and approved the final manuscript.

