## Supplementary material for "Proteome-wide Prediction of the Functional Impact of Missense Variants with ProteoCast": all suplementary figures

### Supplemental Tables

| | Number of genes | Number of unique proteoforms* | Number of proteoforms with predicted landscapes | Number of proteoforms with confident predicted landscapes | Number of residues with confident predictions |
|---|---|---|---|---|---|
| **All** | 13,969 | 22,392 (1.6 ± 1.8 per gene) | 22,169 (99%) | 19,421 (86.7%) | 13,943,646 (96.8%) |
| **Subset with AF2-predicted 3D models** | 13,919 | 22,083 | 21,870 (99%) | 19,212 (87%) | 12,588,986 (97.2%) |

**Table S1: Overview of the predictions for the *Drosophila melanogaster* proteome.** FlyBase version 6.44 of the proteome was used. *data retrieved from FlyBase [1].

| Dataset | Number of SNPs | Number of SNPs unique to dataset | Number of SNPswith confident predictions* | Number of proteins | Number of genes |
|---|---|---|---|---|---|
| DGRP | 177,013 | 53,557 | 137,149 | 12,472 | 10,469 |
| DEST2 | 331,341 | 207,886 | 269,766 | 13,227 | 10,879 |
| Lethal | 1,066 | 787 | 1,004 | 464 | 456 |
| Hypomorphic | 403 | 151 | 148 | 106 | 106 |

**Table S2: Overview of the custom benchmark used to evaluate ProteoCast predictions in the context of organismal fitness.** The benchmark includes four datasets of missense mutations in Drosophila melanogaster: (1) inbred population polymorphisms derived from the DGRP [2]; (2) natural population polymorphisms derived from the DEST2 [3]; (3) ethyl methanesulfonate(EMS)-induced point mutations resulting in lethality, as annotated in FlyBase [1]; and (4) point mutations associated with hypomorphic alleles in FlyBase, which exhibit reduced gene function. *Mutations found in the Lethal dataset and also in DEST2 or DGRP are excluded to ensure a fairer calculation of performance metrics. Additionally, all mutations from the Lethal dataset are excluded from the Hypomorphic dataset.

| Method | Specificity | | Recall | Balanced accuracy | | |
|---|---|---|---|---|---|---|
| | DGRP | DEST2 | Lethal | DGRP | DEST2 | All |
| ProteoCast classification | 0.88 | 0.82 | 0.85 | 0.86 | 0.83 | 0.83 |
| Fixed threshold on predicted scores: -2.4 | 0.87 | 0.81 | 0.81 | 0.84 | 0.81 | 0.81 |

**Table S3: ProteoCast performance on the custom benchmark (DEST2, DGRP, Lethal)**. Balanced accuracy is computed as the average of the specificity (proportion of population variants classified as neutral) and recall (proportion of lethal mutations classified as impactful or mild). We compare ProteoCast classification to binary classifications obtained by applying a universal threshold to the predicted raw scores.

| PDB ID | UniProt ID | Organism | Identity (%) | Coverage (%) | E-value | Publication |
|---|---|---|---|---|---|---|
| 4YUB | Q6XQN6 | *Homo sapiens* | 47.77 | 79 | 2e-86 | Marletta et *al.*[4] |
| 2F7F | Q830Y8 | *Enterococcus faecalis V583* | 44.66 | 67 | 3e-48 | None |
| 1YTD, 1YTK, 1YTE | Q9HJ28 | *Thermoplasma acidophilum* | 37.80 | 18 | 7e-14 | Shin et *al.*[5] |
| 2I14 | Q8TZS9 | *Pyrococcus furiosus* | 27.45 | 60 | 6e-10 | None |
| 1VLP | P39683 | *Saccharomyces cerevisiae* | 22.36 | 32 | 0.002 | Chappie et *al.*[6] |

**Table S4: Homology search for Naprt.** The Naprt-PH proteoform sequence was queried against the Protein Data Bank [7] using PSI-BLAST (2 iterations) [8].

| | **Total number of sites*** | **pLDDT $<$ 70** | | | | |
|---|---|---|---|---|---|---|
| | | **Number of sites** | **Number of sites classified as sensitive** | | **Number of sites standing out from their background** | |
| **PTMs** | 60,428 | 53,737 (89%) | 20,923 (39%) | | 33,866 (63%) | |
| | | | **Overlap (at least 1 res.)** | **Fully included** | **Overlap (at least 1 res.)** | **Fully included** |
| **SLiMs** | 91 | 72 (79%) | 59 (82%) | 34 (47%) | 61 (85%) | 44 (61%) |

**Table S5: Summary table of functional signal retrieved by ProteoCast for post-translational modifications (PTMs) [9] and Short Linear Motifs (SLiMs) [10] in unstructured regions (AlphaFold pLDDT$<$70).** *in proteoforms with 3D structure information.

| Gene | Proteoform | Role | Length in residues | $F_{obs}$ (%) | #Polymorphisms (imp.) | #DEST2 only (imp., mild) | #DGRP only (imp., mild) | #Lethal (neutral) |
|---|---|---|---|---|---|---|---|---|
| Snake | FBpp0082184 | Upstream (protease) | 435 | 93.0 | 28 (0, 7) | 15 (0, 5) | 2 (0, 1) | 0 (0) |
| SPE | FBpp0083832 | Upstream (protease) | 400 | 95.4 | 26 (2, 1) | 14 (2, 1) | 4 (0, 0) | 0 (0) |
| **Spatzle** | **FBpp0084507** | **Ligand** | **326** | **66.5** | **16 (0, 3)** | **8 (0, 2)** | **5 (0, 1)** | **2 (1)** |
| **Toll** | **FBpp0084431** | **Membrane receptor** | **1097** | **76.3** | **54 (0, 3)** | **32 (0, 3)** | **3 (0)** | **9 (2)** |
| Pellino | FBpp0083913 | Intra (U-ligase) | 424 | 64.2 | 7 (3, 0) | 5 (3, 0) | 2 (0,0) | 0 (0) |
| Myd88 | FBpp0087679 | Intra (adaptor) | 537 | 49.9 | 28 (2, 1) | 21 (2, 1) | 3 (0, 0) | 1 (0) |
| Tube | FBpp0291542 | Intra (adaptor) | 462 | 33.3 | 20 (3, 4) | 12 (2, 3) | 3 (0, 1) | 2 (1) |
| Pelle | FBpp0084549 | Intra (kinase) | 501 | 86.5 | 23 (1, 2) | 15 (1, 1) | 4 (0, 1) | 8 (3) |
| Cactus | FBpp0080402 | Intra (inhibitory) | 500 | 72.2 | 20 (1, 2) | 14 (1, 2) | 4 (0, 0) | 0 (0) |
| **Dif** | **FBpp0080561** | **Intra (target TF)** | **667** | **80.1** | **40 (0, 3)** | **19 (0, 2)** | **3 (0, 1)** | **0 (0)** |
| **Dorsal** | **FBpp0080558** | **Intra (target TF)** | **677** | **79.5** | **20 (1, 0)** | **8 (0, 0)** | **7 (0, 0)** | **6 (1)** |
| DEAF-1 | FBpp0074651 | Intra (TF) | 576 | 67.8 | 13 (1, 2) | 10 (1, 1) | 3 (0, 1) | 1 (0) |
| Drs | FBpp0072935 | Downstream | 70 | 81.2 | 1 (0, 0) | 0 (0, 0) | 1 (0, 0) | 0 (0) |
| Mtk | FBpp0086518 | Downstream | 52 | 6.3 | 5 (1, 0) | 2 (1, 0) | 1 (0, 0) | 0 (0) |
| Def | FBpp0087518 | Downstream | 92 | 64 | 10 (0, 1) | 4 (0, 0) | 0 (0, 0) | 0 (0) |
| CecC | FBpp0084980 | Downstream | 63 | 36.6 | 5 (0, 2) | 3 (0, 2) | 0 (0, 0) | 0 (0) |
| AttA | FBpp0086567 | Downstream | 221 | 41.7 | 18 (0, 1) | 5 (0, 1) | 4 (0, 0) | 0 (0) |
| Dipt | FBpp0085802 | Downstream | 106 | 43.5 | 10 (1, 0) | 4 (1, 0) | 0 (0, 0) | 0 (0) |

**Table S6: Overview of the polymorphisms and referenced lethal mutations in 18 proteins from the Toll pathway.** The main components, namely the Toll receptor and its ligands, and the target transcription factors (TF), are highlighted in bold. $F_{obs}$ is the fraction of observed mutations. In the column names, imp., mild, and neutral denote variants classified as impactful, mild or neutral, respectively.

| Number of proteins | Number of variants (% of the full ClinVar set) | | Sensitivity | Specificity | AUC-ROC |
|---|---|---|---|---|---|
| **w confident ProteoCast predictions** | | | | | |
| 2,525 | 61,860 (99%) | ProteoCast | 77.1 | 86.9 | 0.90 |
| **w EVE predictions** | | | | | |
| 1,736 | 41,282 (66%) | ProteoCast | 78.3 | 85.2 | 0.89 |
| | | EVE | 69.0 | 72.5 | 0.89 |
| **w/o EVE *uncertain* class** | | | | | |
| 1,728 | 33,274 (53%) | ProteoCast | 81.9 | 88.4 | 0.92 |
| | | EVE | 86.6 | 88.6 | 0.92 |

**Table S7: Performance of ProteoCast and EVE on the ClinVar dataset.**The table reports the number of proteins, evaluated mutations, sensitivity, specificity, and AUC-ROC for different evaluation settings. Percentages in parentheses indicate the proportion of mutations relative to the full ClinVar set (62,727 mutations across 2,525 proteins). For Proteocast, mutations classified as neutral are predicted as benign and the rest as pathogenic. EVE classification uses the same categories as ClinVar.

**Table S8: phylo-HMM dataset.** Curated set of 500 phosphorylation sites, localization signals, degradation signals, SUMO sites, and interaction motifs in *Saccharomyces cerevisiae* defined in Nguyen Ba et *al.* (2012) [11].

| Number of sites (% of all 521 sites) | Number (%) detected by phylo-HMM | Number (%) within segments having elevated sensitivity | |
|---|---|---|---|
| | | Overlap (at least 1 residue) | Fully included |
| **pLDDT<70** | | | |
| 437 (84%) | 99 (23%) | 240 (55%) | 186 (43%) |
| **phylo-HMM disorder** | | | |
| 347 (67%) | 104 (30%) | 175 (50%) | 132 (38%) |
| **phylo-HMM disorder & pLDDT<70** | | | |
| 333 (64%) | 99 (30%) | 169 (51%) | 127 (38%) |

**Table S9: Comparison of ProteoCast and phylo-HMM in detecting functional motifs in *S. cerevisiae* proteins.** The full set is the one described in [11] except 5 for which the input MSA was of too low quality. The "phylo-HMM disorder" set was identified through a multi-step disorder prediction pipeline in Nguyen Ba et *al.* (2012).

| Number of sites | Number of protein sequences | Number (%) within segments having elevated sensitivity | | |
|---|---|---|---|---|
| | | Overlap (at least 1 residue) | Overlap (at least 50%) | Fully included |
| 62 | 50 | 53 (85%) | 40 (65%) | 25 (40%) |

**Table S10: ProteoCast detection of binding sites annotated in CAID3 [12].**

| Predictor | AUC | APS | F1 max | bac |
|---|---|---|---|---|
| bindEmbed21IDR-rawGeneral | 0.641 | 0.514 | 0.589 | 0.604 |
| bindEmbed21IDR-idrGeneral | 0.635 | 0.497 | 0.587 | 0.602 |
| LIPNet | 0.619 | 0.507 | 0.558 | 0.592 |
| ProBiPred-protein | 0.613 | 0.472 | 0.563 | 0.587 |
| AlUPred-2-binding | 0.590 | 0.470 | 0.557 | 0.577 |
| **ProteoCast-delta-mu_segment** | **0.587** | **0.455** | **0.556** | **0.600** |
| bindEmbed21IDR-rawNuc | 0.582 | 0.467 | 0.557 | 0.575 |
| bindEmbed21IDR-idrNuc | 0.579 | 0.460 | 0.557 | 0.573 |
| DisoRDPbind-rna | 0.575 | 0.454 | 0.561 | 0.554 |
| **ProteoCast-mu_residue** | **0.572** | **0.448** | **0.561** | **0.555** |
| MoRFchibi-light | 0.564 | 0.423 | 0.560 | 0.561 |
| MoRFchibi-web | 0.548 | 0.419 | 0.558 | 0.541 |

**Table S11: AUC, average precision score (APS), maximum F1-score, and balanced accuracy (bac) computed at the residue level for the CAID3 dataset [12].** The set of positives comprises the residues labelled as binding sites and the set of negatives comprises all other residues labelled as disordered – corresponding to the "Binding-IDR" CAID3 challenge. We retained the 10 best-performing methods and ordered them according to the AUC. ProteoCast comes in two flavours: **ProteoCast-delta-mu_segment**, where we use the maximum difference in mutational sensitivity between the considered segment and its two neighbouring segments as the predictive metric, and **ProteoCast-mu_residue**, where we use per-residue mutational sensitivity as the predictive metric.

### Supplemental Figures

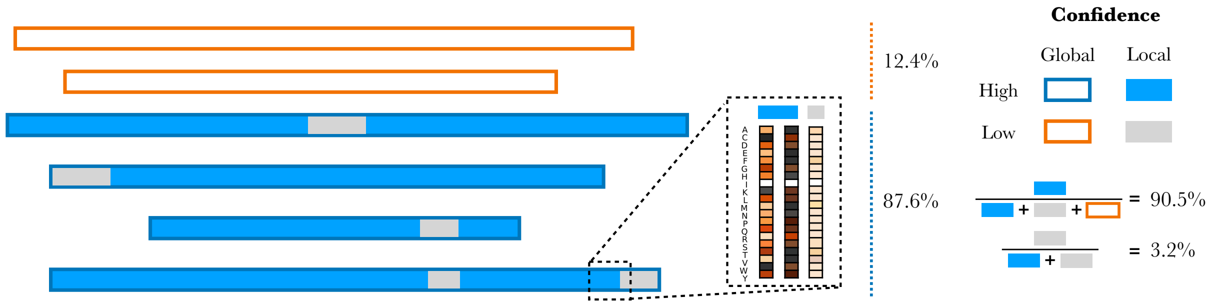


**Figure S1: Overview of global and local confidence metrics.** The rectangular outlines (orange or blue) indicate global confidence while the internal coloring (blue or grey) indicates local confidence (Percentages based on n = 22,169 proteoforms analyzed). The insert illustrates archetypal scenarios for high and low residue confidence. The left column has high resolution, meaning that ProteoCast can distinguish the effects of the different substitutions at that protein position. By contrast, the two other columns show a narrow distribution of scores. In the middle, all substitutions are predicted to induce strong effects, implying high evolutionary conservation of the wild-type amino acid. On the right however, the low-resolution and weak predicted values may indicate that the amount of input evolutionary information is not sufficient to make a reliable prediction.

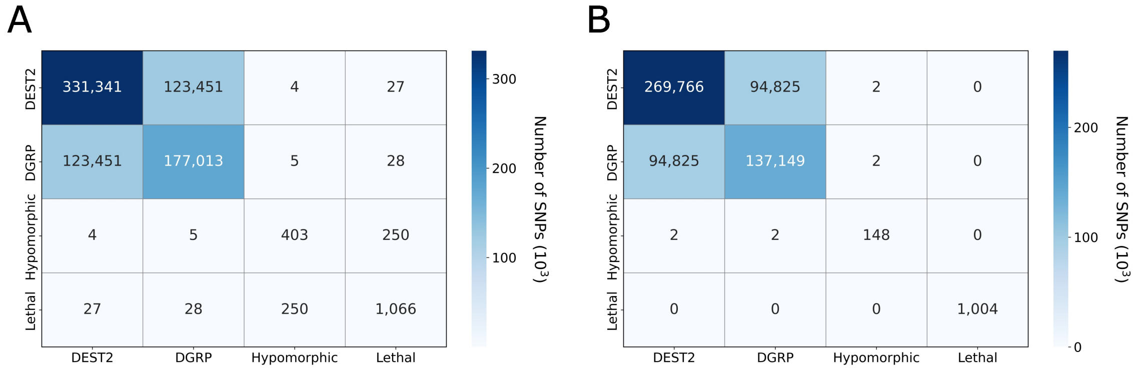


**Figure S2: Overview of the custom benchmark for evaluating ProteoCast predictions in the context of organismal fitness. A.** All mutations. **B.** Mutations whose effect is predicted with high global and local confidence and that are not annotated as both lethal and observed in fly populations (DEST2 or DGRP). Additionally, mutations referenced as lethal are excluded from the Hypomorphic dataset.

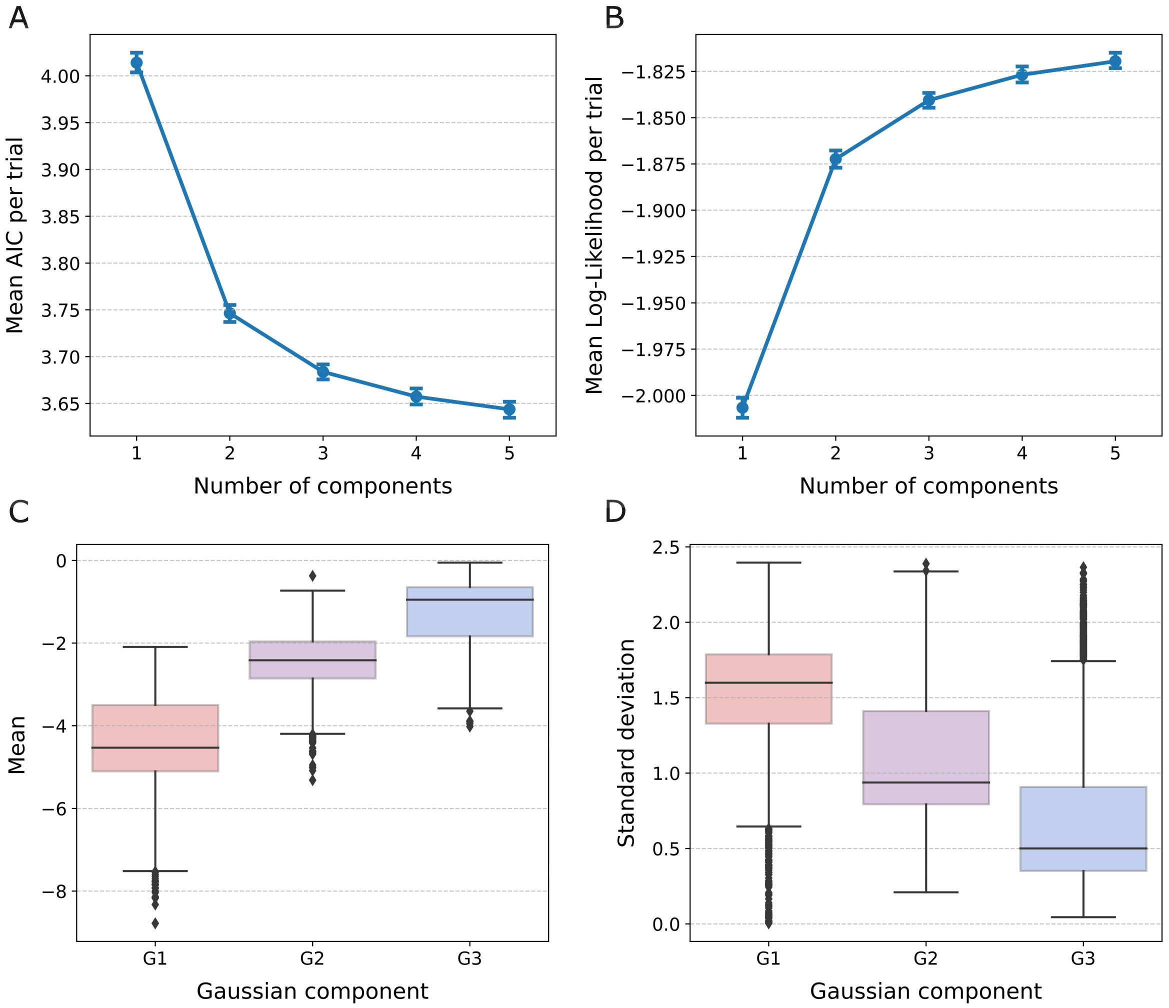


**Figure S3: Gaussian Mixture Model (GMM) fit and component summary statistics. A-B.** Evaluation of GMM fits to mutational landscape predictions across 20 trials of 100 randomly selected proteoforms with 1-5 components (error bars: 95% CI across n=20 trials). **A.** Mean Akaike Information Criterion (AIC) per trial. **B.** Mean log-likelihood per trial. **C-D.** Distributions of the estimated means (C) and standard deviations (D) for the three Gaussian components across the same set of proteoforms.

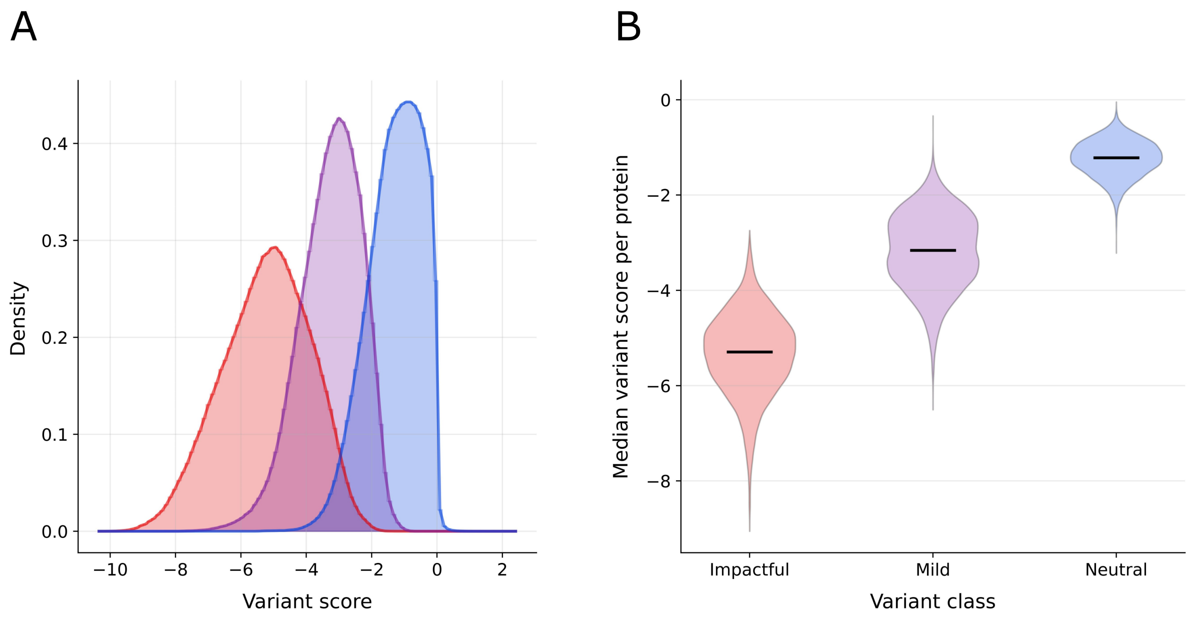


**Figure S4: Distribution of predicted variant effect scores across variant classes. A.**Density distributions of variant scores for all variants, stratified by predicted variant class: impactful (red), mild (purple), and neutral (blue). **B.** Distribution of median values across proteins for each variant class.

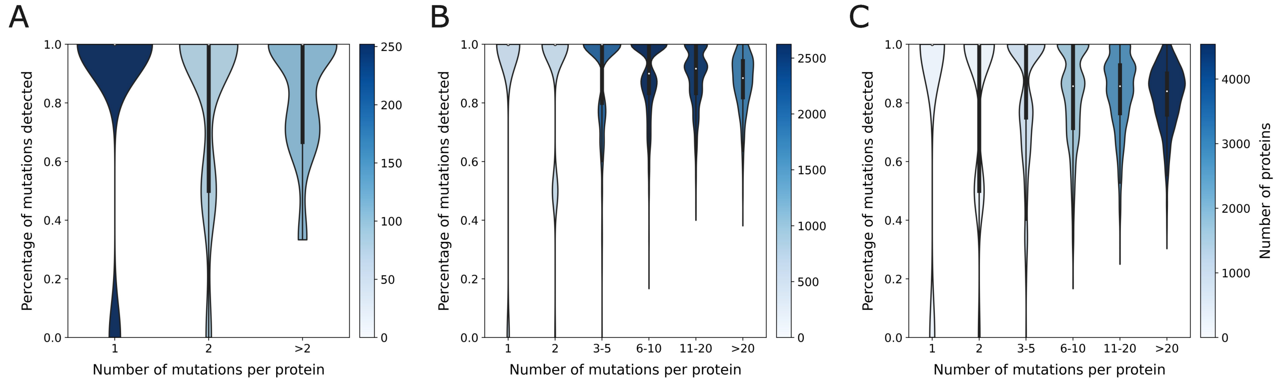


**Figure S5: Distribution proportion of mutations correctly classified for individual proteins.** Proteins are grouped by the number of annotated mutations in our benchmark: 1, 2, [3-5], etc. mutations per protein. (A) lethal mutations (n=252, n=88, n=116 proteins in each group), (B) DGRP variants (n=703, n=727, n=2082, n=2624, n=2514, n=1819 proteins in each group), (C) DEST2 variants (n=306, n=326, n=1079, n=1774, n=2855, n=4539 proteins in each group). The colour scales indicate the number of proteins within each group.

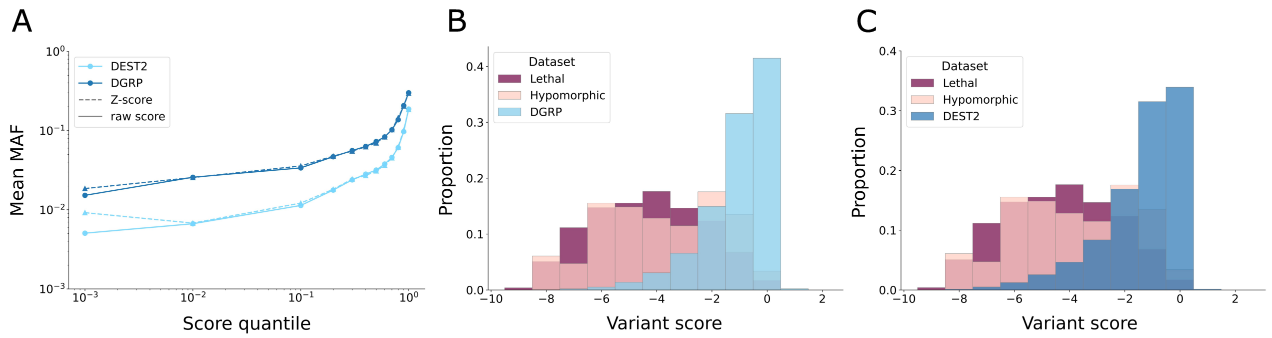


**Figure S6: Distribution of scores for the different types of mutations in our benchmark. A.** Minor Allele Frequencies (MAF) in function of predicted raw- and Z-scores for inbred and natural polymorphisms (n=177013 DGRP variants, n=331341 DEST2 variants). We report the MAF mean values in y-axis for different raw- and Z-score quantile bins $[0, 10^{-3})$, $(10^{-3}, 10^{-2}]$, ..., $(10^{-1}, 1]$ in x-axis. **B-C.** Distributions of predicted raw scores for the four missense mutation subsets in our custom benchmark: lethal (deep purple, B-C), hypomorphic alleles (pale pink, B-C), DGRP (sky blue, B), and DEST2 (steel blue, C). Sample sizes (n) for each dataset are provided in Table S2.

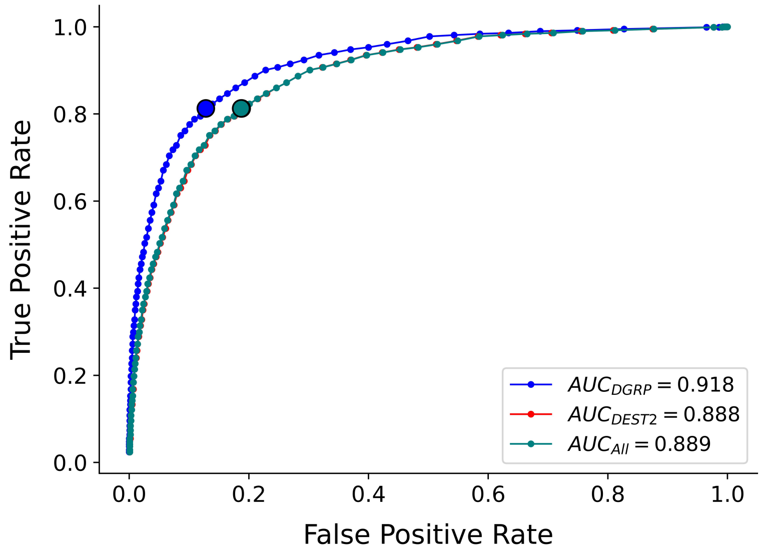


**Figure S7: Receiver Operating Characteristic (ROC) curves comparing predictive performance based on ProteoCast raw scores.** The True Positive Rate (sensitivity) is plotted against the False Positive Rate for three datasets differing in their selection of negative examples: DGRP-only (blue), DEST2-only (red), and All (teal), which includes the overlap between DEST2 and DGRP. The Area Under the Curve (AUC) values are provided in the legend. Large markers indicate thresholds with optimal performance. The DEST2 and All datasets exhibit nearly identical performance.

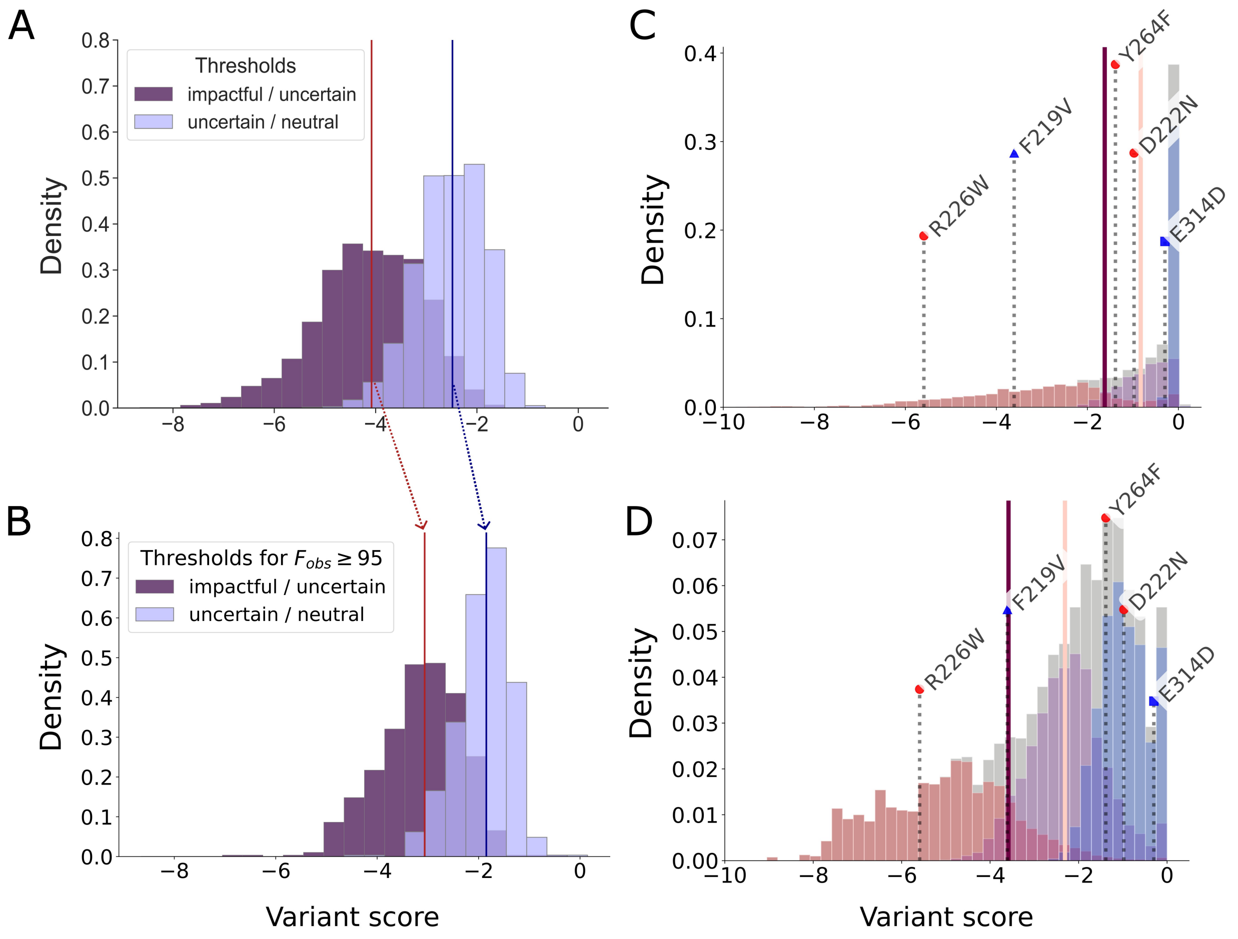


**Figure S8: A-B. Shift in GMM-based classification thresholds upon increasing the fraction of observed mutations. A.** Entire proteome. The median values for the impactful-mild and mild-neutral thresholds are of -4.08 and -2.48, respectively. **B.** Proteins with nearly complete coverage of their mutational landscape in the alignment ($F_{obs} \geq 95$). The median values increase up to -3.06 and -1.85. **C-D. ProteoCast classification for the protein-serine/threonine phosphatase Flapwing mutations (flw-PA proteoform, FBpp0071381).** The five known mutations are highlighted on the raw score distributions of Flapwing itself (C) and of Yorkie (D). The three Gaussians fitted to each distribution are colored in red, purple, and blue respectively. The pale pink vertical line represents the threshold above which mutations are considered *neutral*, the deep purple vertical line marks the threshold below which mutations are considered as *impactful*. Mutations with scores between the two threshold values are considered *mild*. Two lethal mutations that would appear as neutral in Yorkie (D) are classified as mild in Flapwing (C) highlighting the need for adaptive thresholds for variant classification.

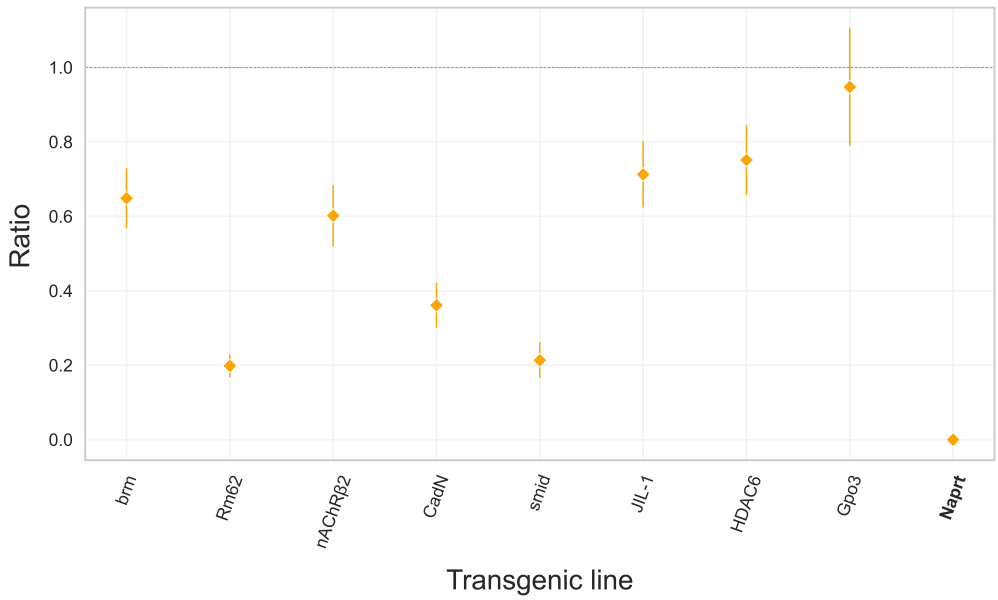


**Figure S9: Developmental lethality associated with gene-specific RNAi knockdowns.** Ratio of *da::*GS>UAS-RNAi individuals developing out of two conditions RU0 ($\mu$g/mL of RU486) - negative control, the RNAi is not induced -, and RU20 - the RNAi is expressed at the maximum minducer concentration for that system. We present here the ratio of the total number of adults eclosing from the induced condition divided by the total number of adults eclosing from the control condition. n=5 independent vials per condition (RU0 and RU20). Error bars represent SEM from n=5 independent biological replicates (matched vials). List of genes targeted with RNAi in the experiment: *nAChRβ2* (FBgn0004118), *CadN* (FBgn0015609), ***Naprt*** (FBgn0031589), *Gpo3* (FBgn0028848), *JIL-1* (FBgn0020412), *mrj* (FBgn0034091), *smid* (FBgn0016983), *Rm62* (FBgn0003261).

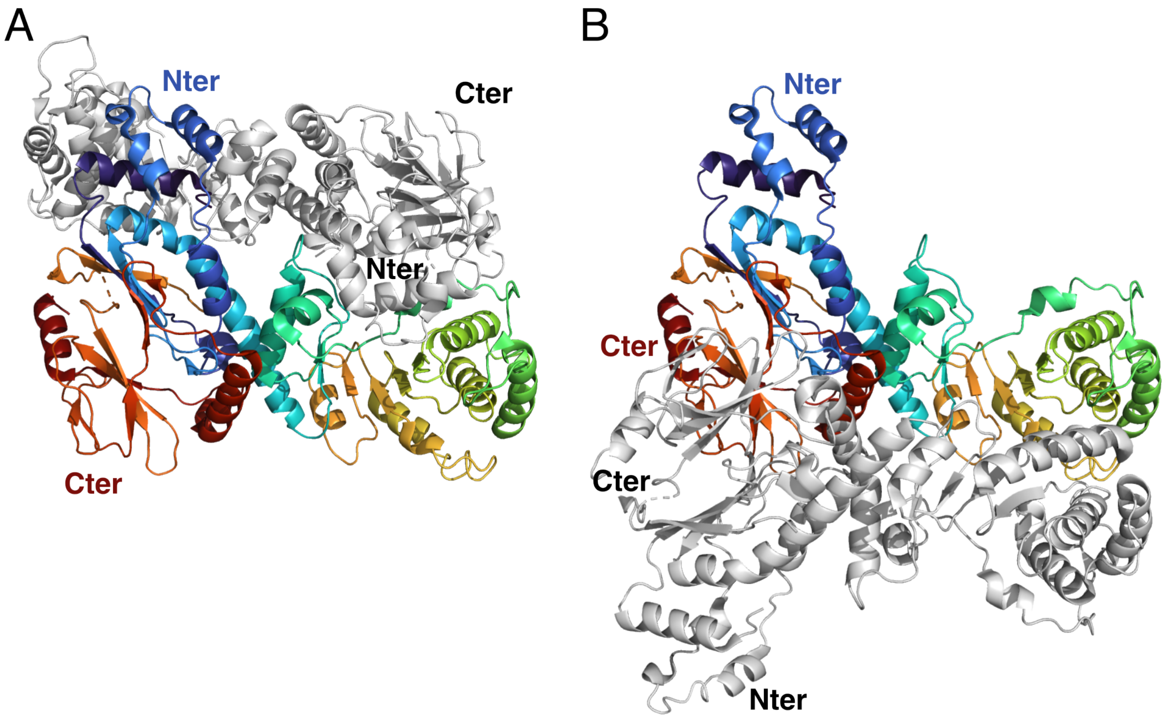


**Figure S10: Human NAPRT homo-dimer.** The coordinates were retrieved from the PDB entry 4YUB. One monomer is taken as reference and colored in rainbow according to the residue index in the protein sequence. The second monomer is colored in white. **A.** Biologically relevant head-to-tail homo-dimeric arrangement of human NAPRT described in Marletta and co-authors (see Fig. 2 in [4]). **B.** Head-to-head dimeric arrangement incorrectly defined as the biological assembly and asymmetric unit in the Protein Data Bank.

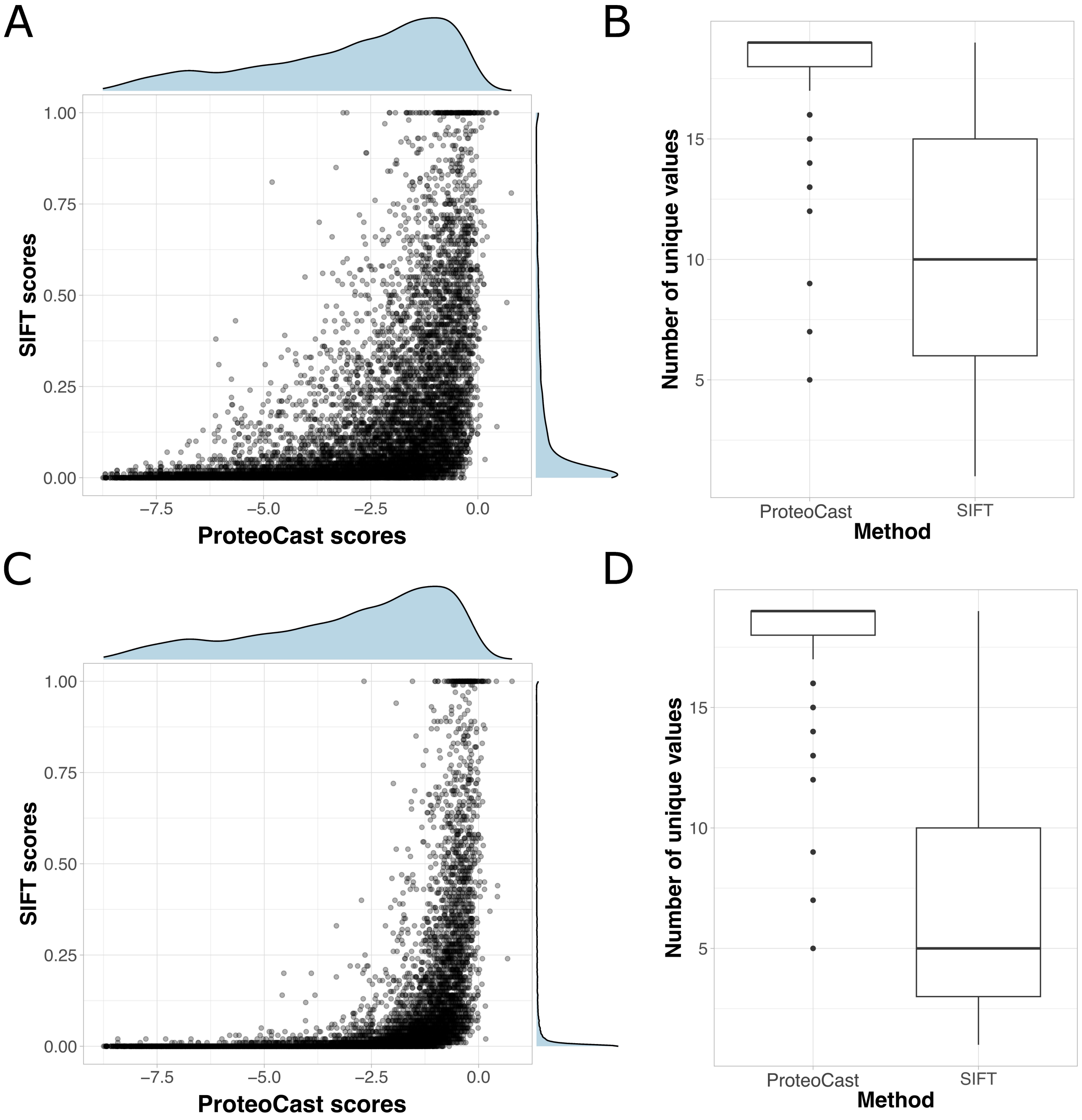


**Figure S11: Comparison between ProteoCast and SIFT for Naprt-PH.** We ran SIFT through the webserver `https://sift.bii.a-star.edu.sg/` in single-sequence mode (A-B) or inputting the same alignment as that used for ProteoCast (C-D). **A,C.** SIFT scores in function of ProteoCast raw scores. The respective distributions are shown on top and on the right. SIFT scores below 0.05 indicate impactful variants. **B,D.** Distributions of the number of unique values per residue for the two methods. In all plots, the values obtained for the poorly conserved unstructured regions 1-21 and 301-407 are excluded to reduce the risk of bias they might introduce, n=10,146 possible variants for Naprt-PH". The full ProteoCast predictions for Naprt-PH can be accessed at: `https://proteocast.ijm.fr/results/results/?q=FBpp0306840`.

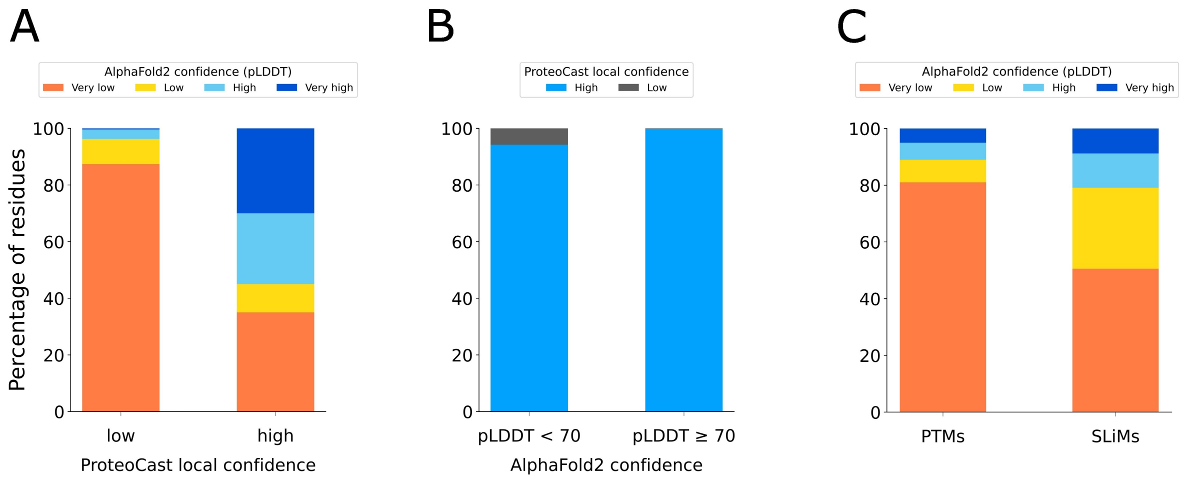


**Figure S12: Relationship between AlphaFold2 pLDDT confidence and ProteoCast local confidence. A.** Proportion of residues in different AlphaFold2 pLDDT categories (very low, low, high, very high), reflecting confidence in their predicted 3D coordinates, shown separately for low (n=361,895 residues) and high (n=12,588,986 residues) ProteoCast local confidence. **B.** Proportion of residues classified as high or low confidence by ProteoCast, grouped by AlphaFold2 pLDDT confidence thresholds (pLDDT $< 70$ vs. pLDDT $\geq 70$, n=6,023,625 and n=6,927,256 ). **C.** Distribution of AlphaFold2 pLDDT scores for PTMs and SLiMs. In total, there are $\sim$60K PTMs and 91 SLiMs.

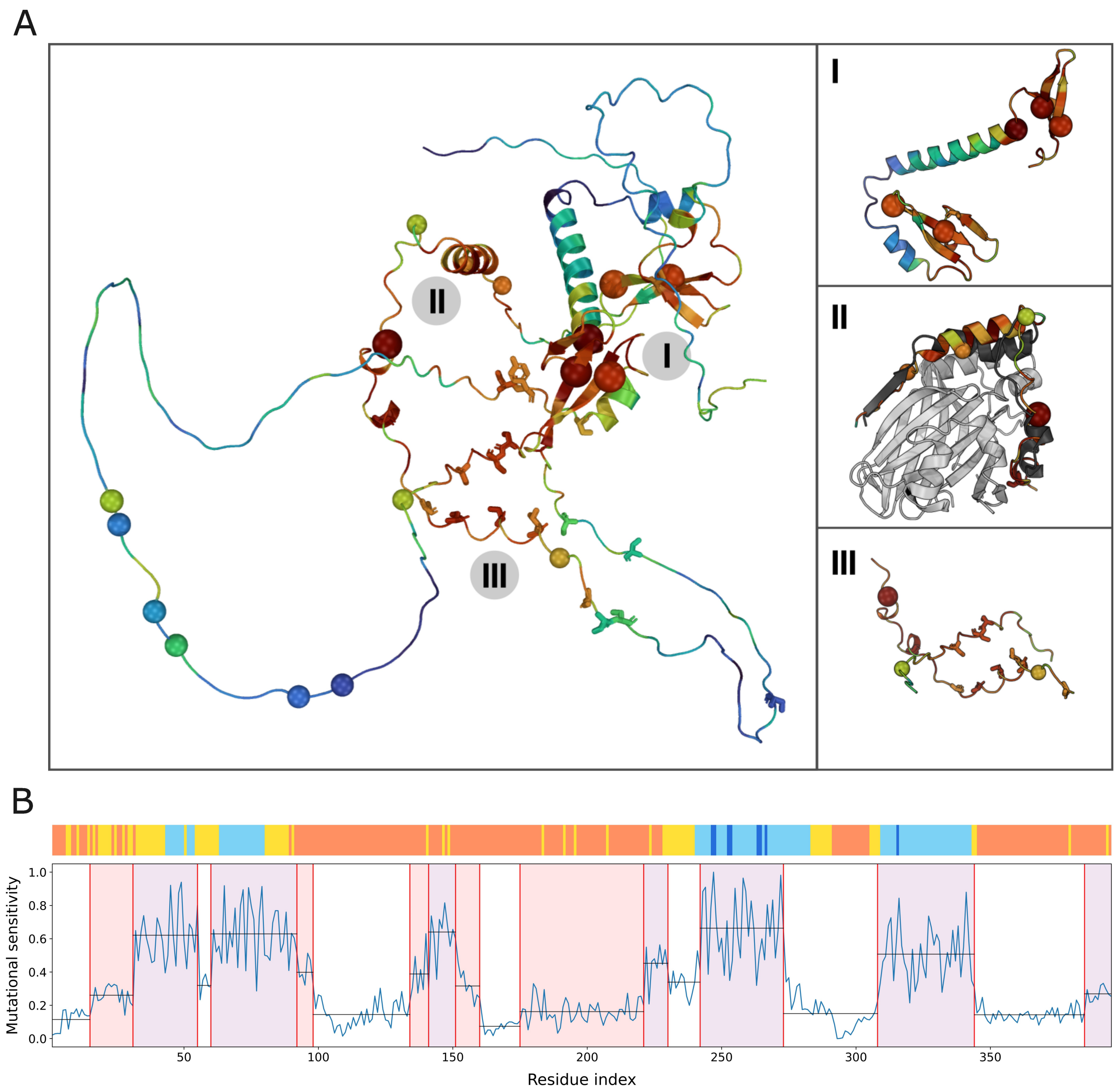


**Figure S13: Mutational sensitivity in Yorkie protein. A.** Per-residue mutational sensitivity scores mapped onto the AlphaFold2-predicted 3D model (*yki* gene, FlyBase proteoform id: FBpp0301274, Uniprot id: Q45VV3). The colour gradient goes from blue for highly tolerant positions to red for highly sensitive residues. Annotated lethal mutations are highlighted with big spheres while natural and inbred polymorphisms are depicted with small spheres. The known PTM sites are shown as sticks. Three closeup views are shown on the right. I. Region from residue 241 to 343 encompassing the two annotated WW domains. II. Region from residue 31 to 80 sharing similarity with the human YAP1 segment interacting with TEAD. The experimental structure of the YAP1-TEAD complex (PDB id: 3KYS) is superimposed, with YAP1 in dark grey and TEAD in light grey. III. Segments 61-98 and 135-160 identified by ProteoCast as standing out from their surrounding background. **B.** AlphaFold pLDDT (top) and segmented mutational sensitivity profile (bottom). **Top.** Orange and yellow indicate very low to low pLDDT, while light and dark blue represent high to very high pLDDT regions. **Bottom.** The identified segments are coloured red if their mean value (horizontal line) is higher than one neighboring segment, and purple if it is higher than both neighbouring segments.

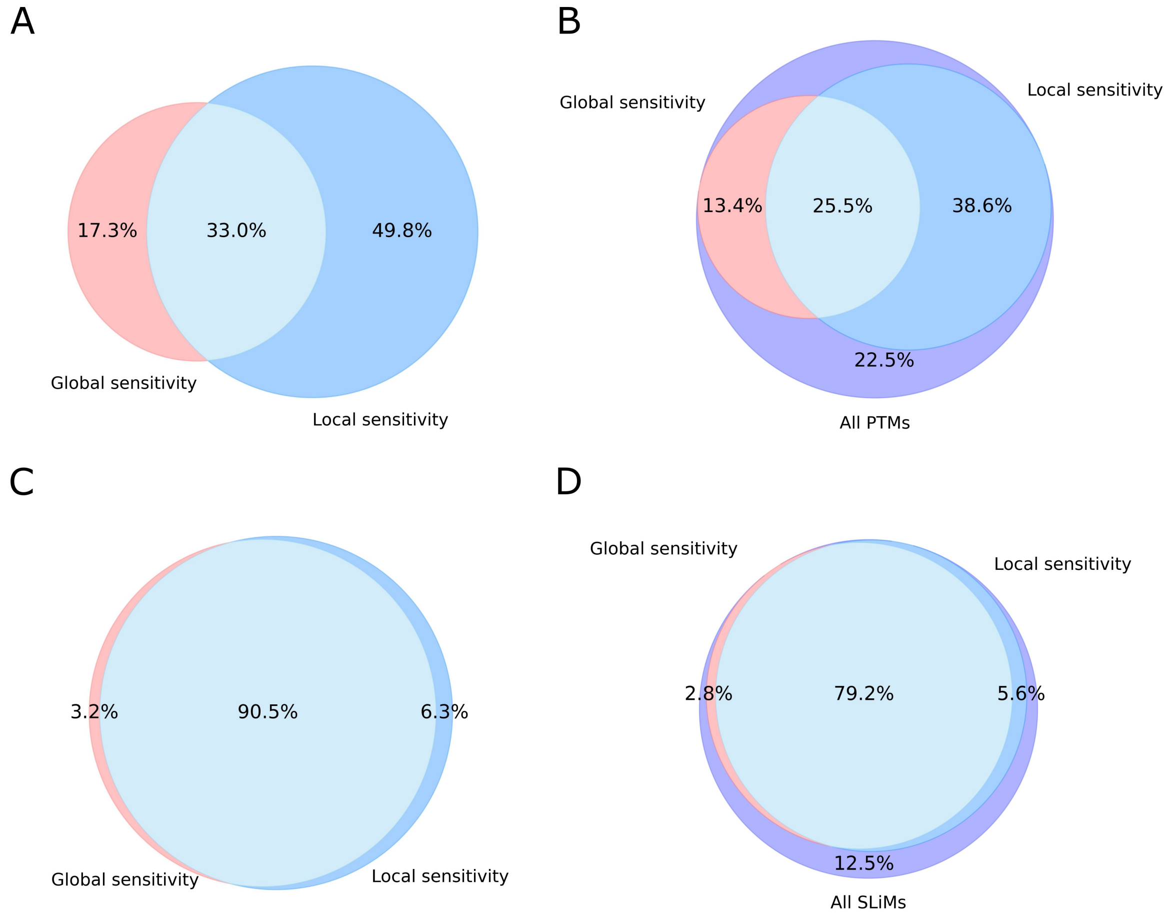


**Figure S14: ProteoCast local and global mutational sensitivity-based detection of post-translational modification sites (A-B) and short linear motifs (C-D).** We focus on regions with low AlphaFold2 pLDDT score (below 70). Glocal sensitivity refers to ProteoCast residue classification where sensitive residues have less than 10 neutral substitutions. Local sensitivity refers to segments whose mutational sensitivity stands out from their surroundings. The Venn diagrams show the complementarity between the two approaches. **A-B.** Detection of post-translational modification (PTM) sites, considering the subset of detected sites (A) or all sites (B). **C-D.** Detection of short linear motifs (SLiMS), considering the subset of detected motifs (C) or all motifs (D).

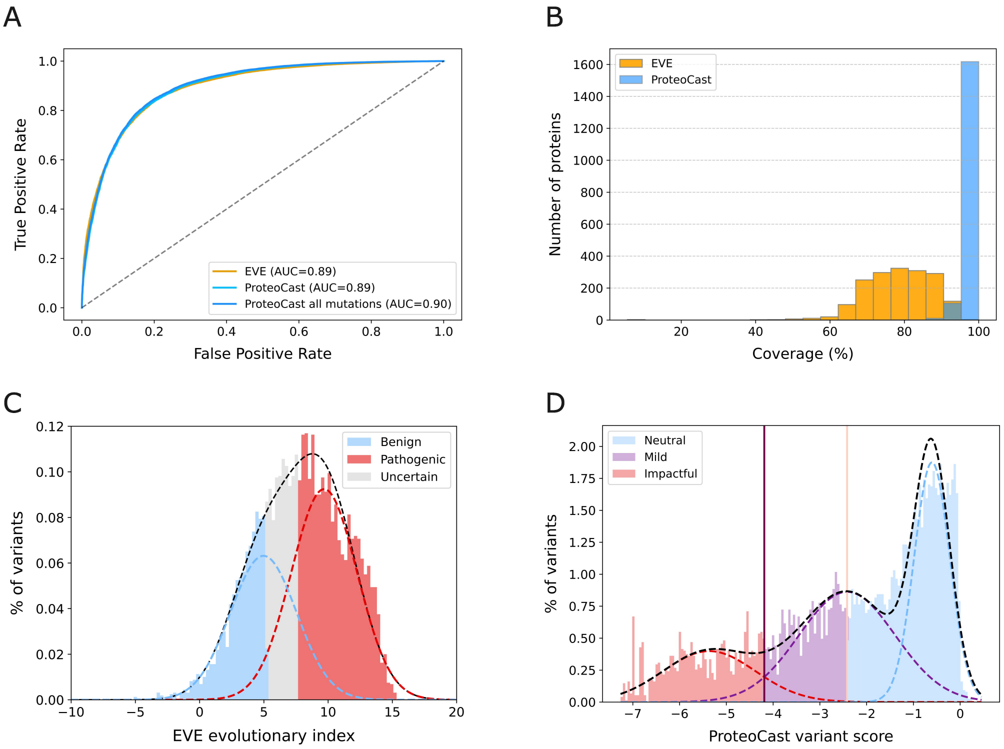


**Figure S15: Comparison of EVE and ProteoCast predictive performance, coverage, and classification procedure on ClinVar. A.** Receiver Operating Characteristic (ROC) curves for ProteoCast (cyan and blue) and EVE (orange) from using the raw predicted scores. The cyan and orange curves correspond to the subset of 41,282 ClinVar mutations where both methods provide predictions, while the blue curve corresponds to the full ClinVar set (61,860 mutations). **B.** Distribution of per-protein mutational landscape coverage for EVE (orange) and ProteoCast (blue), defined as the percentage of possible missense mutations for which a prediction is available. Histograms show that ProteoCast (blue) achieves consistently higher coverage across proteins compared to EVE (orange), with most of the mutational landscape nearly fully covered by ProteoCast. **C-D.** Predicted score distributions for TP53 protein (P53_HUMAN, P04637). The fitted Gaussians (dashed lines) illustrate the classification scheme for EVE (C) and ProteoCast (D). The predicted classes are depicted by colours: benign in blue, pathogenic in red, and mild in grey for EVE (EVE_classes_75_pct_retained_ASM) (C), neutral in blue, likely impactful in purple, and impactful in red for ProteoCast (D). Vertical lines mark decision thresholds between classes for ProteoCast (D).

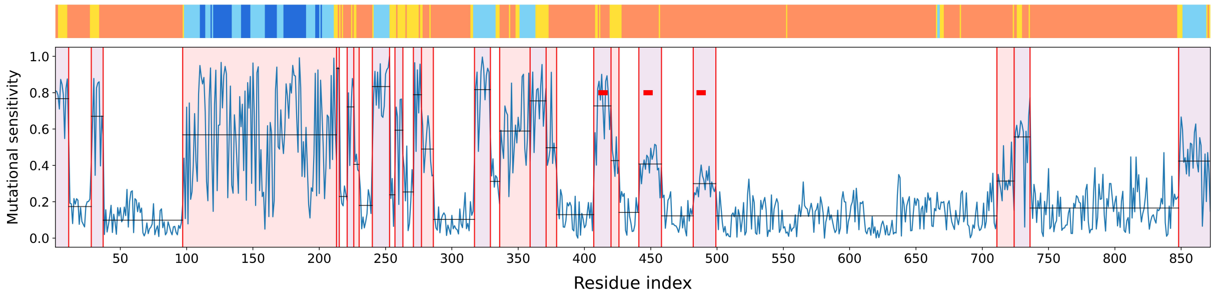


**Figure S16: ProteoCast segmented mutational sensitivity profile for yeast protein SCD5 (YOR329C).** The segments' boundaries are indicated by vertical red lines and their average sensitivity by horizontal black lines. The segments are colored according to whether they have higher sensitivity than both surrounding segments (in purple), or only one neighbour (red), or none (white). Three identical motifs "LKPTATG" known from the literature [13] are highlighted with red squares. All three fall within ProteoCast purple segments while only the first one is detected by phylo-HMM [11]. The pLDDT profile is displayed on top, with orange for pLDDT<50, yellow between 50 and 70, light blue between 70 and 90, and dark blue above 90.

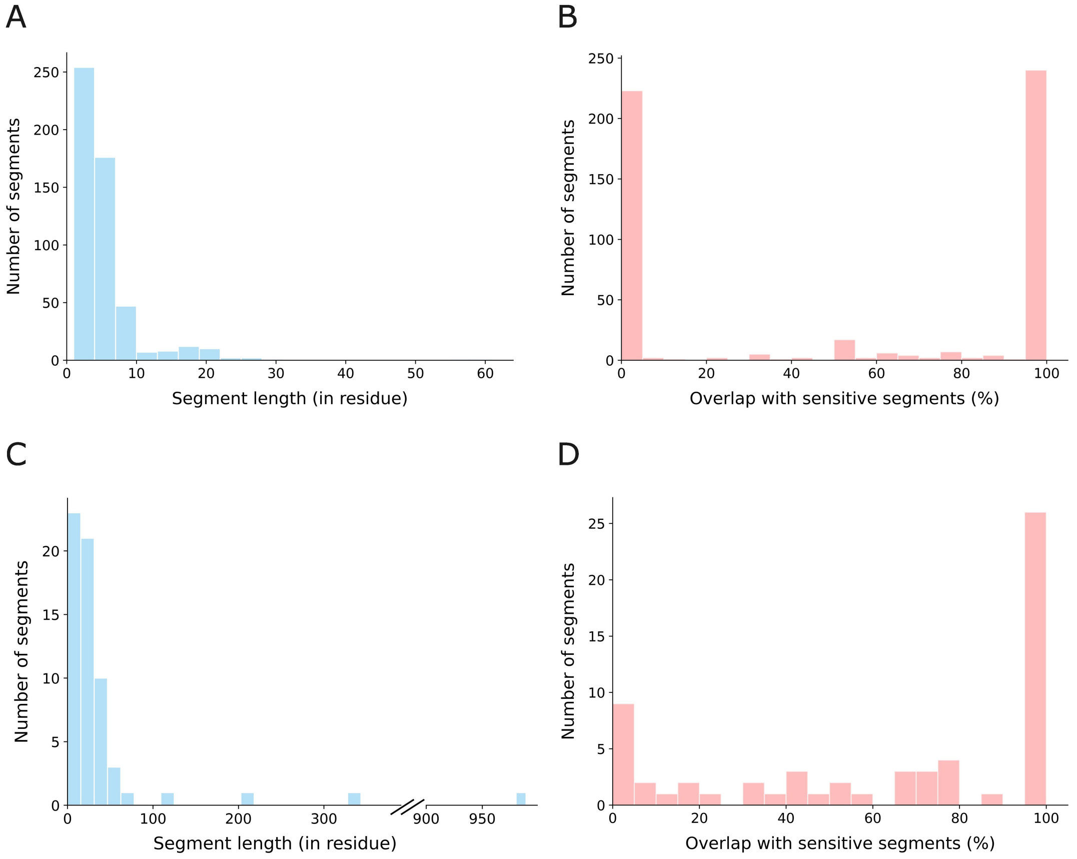


**Figure S17: Length and overlap with sensitive segments for experimentally validated regulatory and binding sites. A-B.** Curated set from Nguyen Ba et *al.* (2012) [11]. **C-D.** Binding-IDR dataset from CAID3 [12]. **A.** Length distribution of curated regulatory sites from Nguyen Ba et *al.* (2012). **A.** Overlap with sensitive segments for the same sites, expressed as the percentage of residues in a site that fall within ProteoCast sensitive segments. **C.** Length distribution of CAID3 binding sites in disordered regions (broken x-axis to display very long sites). **D.** Overlap with sensitive segments for CAID3 binding sites. Overlap with sensitive segments is computed as $\mathbf{O}_{\text{sens}} = \frac{\text{Number of positions within segments class 1 or 2}}{\text{Total number of positions in the site}} \times 100\%$.

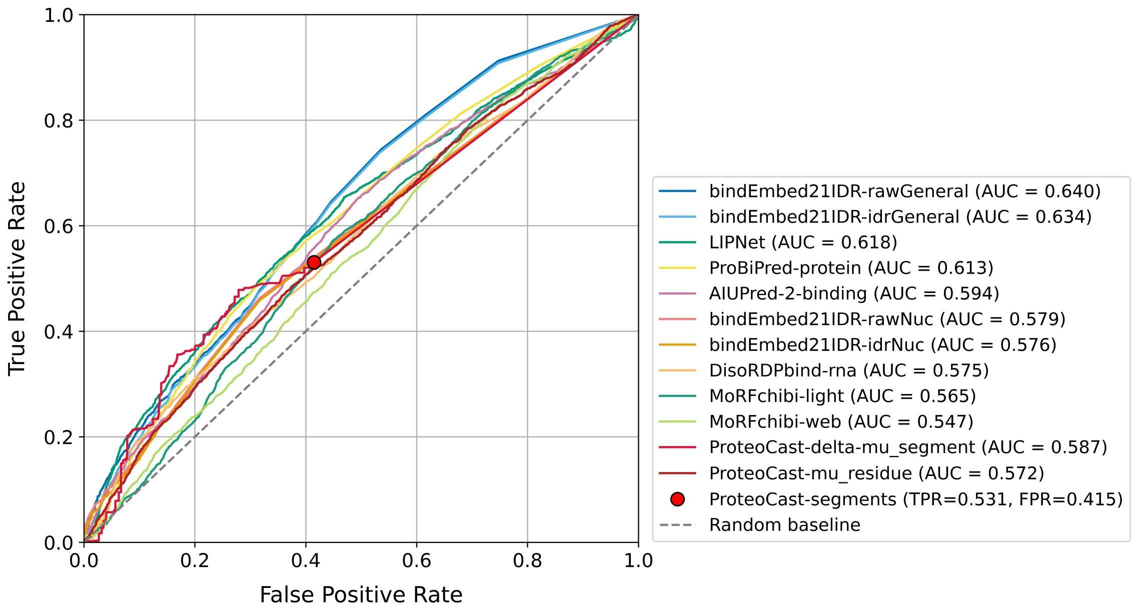


**Figure S18: ROC curves for CAID3 Binding-IDR dataset predictions [12].** We focus on the top 10 CAID3 predictors and ProteoCast. The area under the curve (AUC) is indicated for each method. **ProteoCast-segments**: segments with higher sensitivity than at least one neighbouring segment are considered as positives. **ProteoCast-delta-mu_segment**: we use the maximum difference in mutational sensitivity between the considered segment and its two neighbouring segments as the predictive metric. **ProteoCast-mu_residue**: we use per-residue mutational sensitivity as the predictive metric.

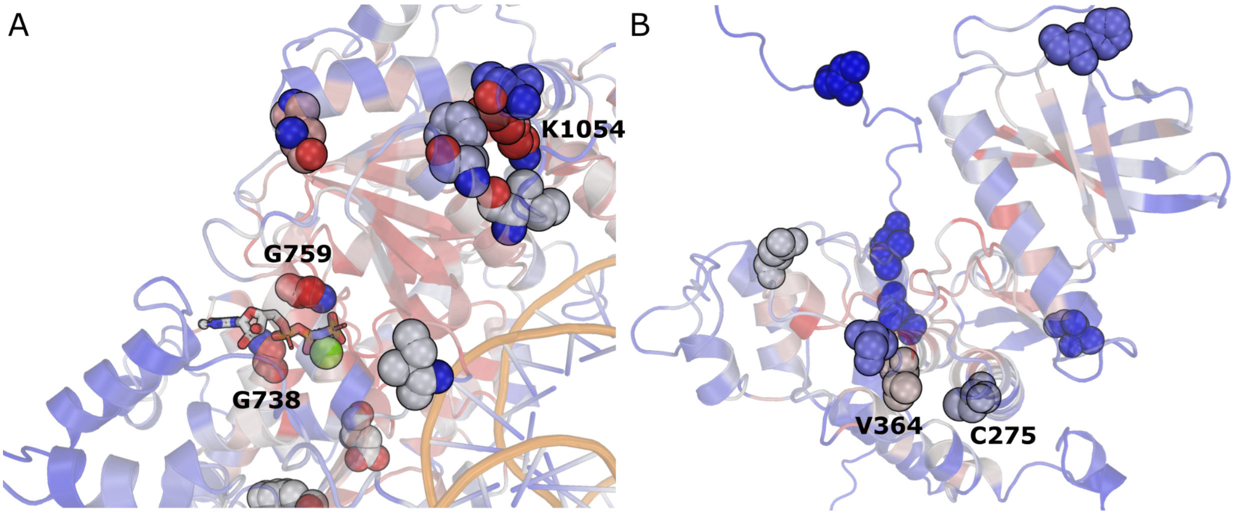


**Figure S19: 3D mapping of natural polymorphism observed in wild flies and predicted as having strong effects on the protein function.** The 3D models for the query proteins were retrieved from the AlphaFold Database or predicted with ColabFold. They are depicted as cartoons and colored according to their per-residue predicted mutational sensitivity. **A.** Close-up view of the putative ATP- and DNA-binding domain of Mi-2 (FBpp0099808), a nuclear ATP-dependent nucleosome remodeler of the CHD family. The ATP and DNA were positioned by superimposing the bound 3D structure of its human homolog CHD4 (PDB id: 6RYR). The location of known mutations predicted as impactful are highlighted with spheres. The lethal mutation sites G738 and G759 are located near ATP while some of the sites exhibiting naturally occurring mutations form a 3D cluster near the DNA-binding interface. The lethal mutations G738D and G759S, and the DEST2 mutations K1054T have the most negative raw scores (below $-6$). **B.** Close-up view of the kinase domain of Doa (FBpp0289026), involved in somatic sex determination. The known mutation sites are highlighted with spheres. The residue V364, whose substitution into G is observed in DEST2 despite being predicted as having a strong effect (raw score $< -6$), is located very close to the lethal mutation site C275.

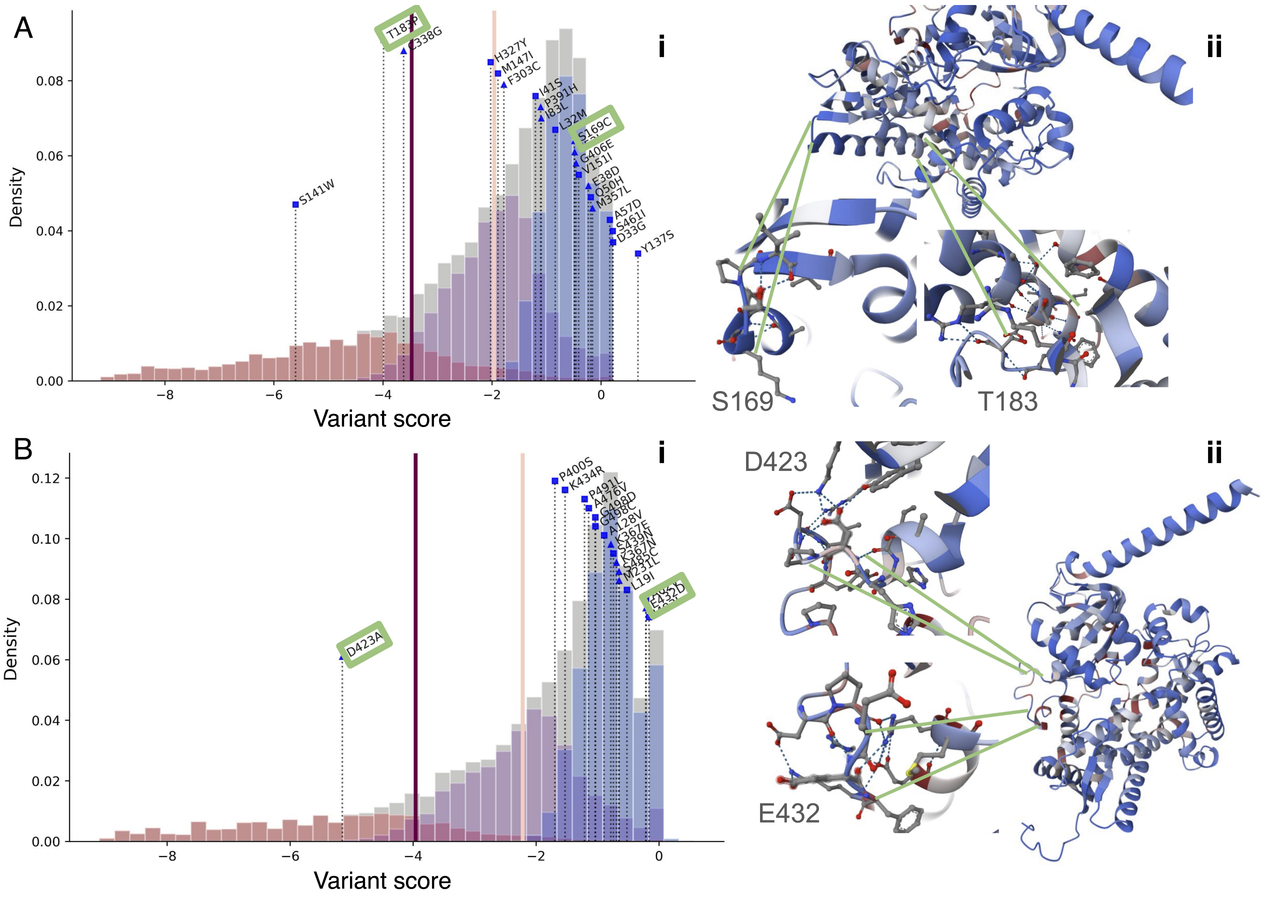


**Figure S20: ProteoCast-informed discrimination between co-segregating natural polymorphisms. Panels i** show the ProteoCast score distributions. Panels ii show the AlphaFold-predicted structures colored with the ProteoCast-predicted mutational sensitivity. Cyp6a9 **(A)** and Cyp317a1 **(B)** belong to a family of detoxifying proteins catabolising insecticides amongst other xenobiotics that have been identified as showing natural polymorphism regionally affected, likely as a function of local agricultural uses. The proximity between the two missense mutation sites highlighted for each protein makes them likely to co-segregate and undistinguishable using a GWAS approach. The ProteoCast predictions can significantly help to prioritise one mutation over another, whether the two mutation sites are located in different secondary structure elements – T183P in an $\alpha$-helix versus S169C in a loop **(Aii)**, or they are both part of an unstructured region and one amino acid substitution implies more drastic physico-chemical changes than the other – D423A more drastic than E432D **(Bii).**

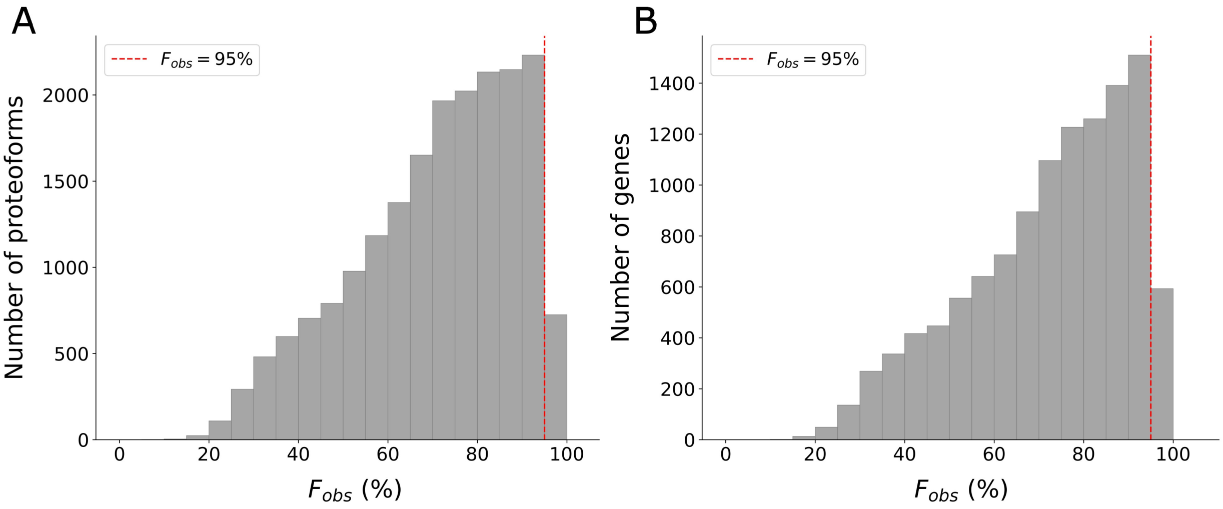


**Figure S21: Distribution of fraction of observed mutations, $\mathbf{F_{obs}}$. A.** By unique proteoform. **B.** By gene, taking the maximum value over the proteoforms of a given gene. We retained only the 19,421 proteoforms, coming from 11,564 genes, with high global confidence predictions.

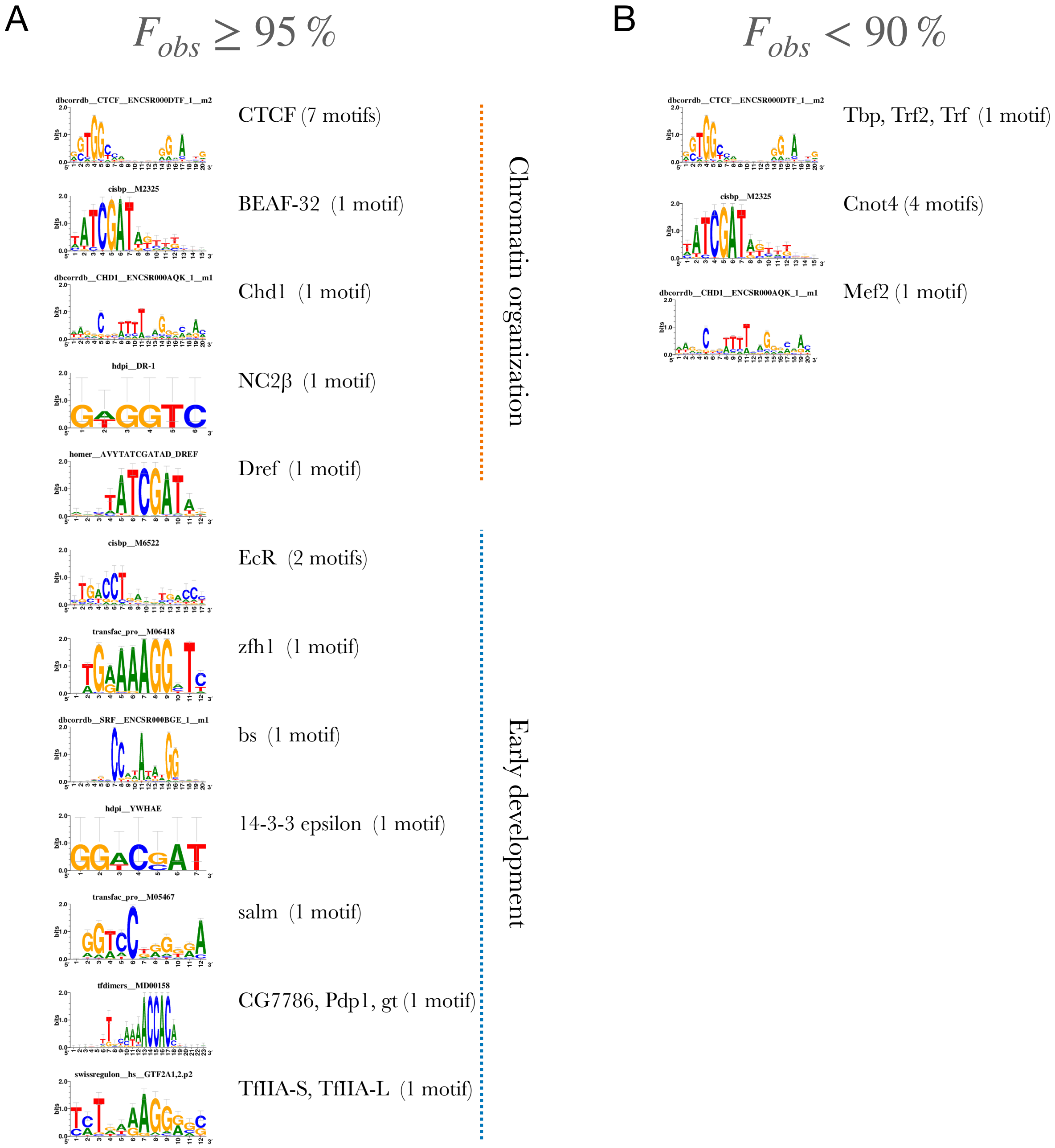


**Figure S22: Motif enrichment in subsets of genes with different fractions of observed mutations. Motif enrichment NES > 5.**

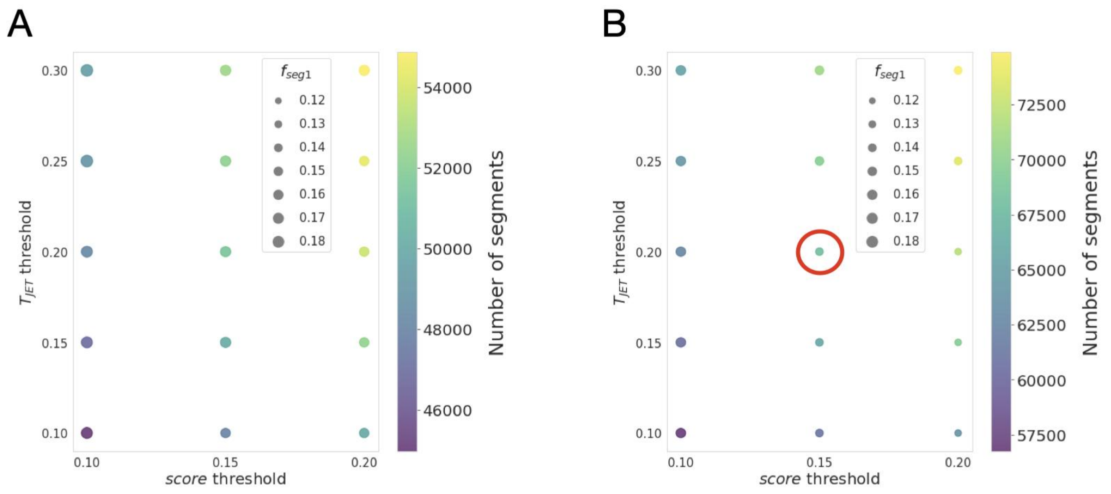


**Figure S23: Assessment of different threshold configurations for detecting residues with low local confidence.** Dot plots illustrate the impact of varying *score* and $T_{JET}$ thresholds on the number of segments with low confidence scores ($N_{seg}$, depicted by a color scale) and the fraction of segments of length one ($f_{seg1}$), with (A) $\sigma_{pred}(s^*, i) = 0.1$ and (B) $\sigma_{pred}(s^*, i) = 0.15$. The red circle highlights the selected threshold combination.

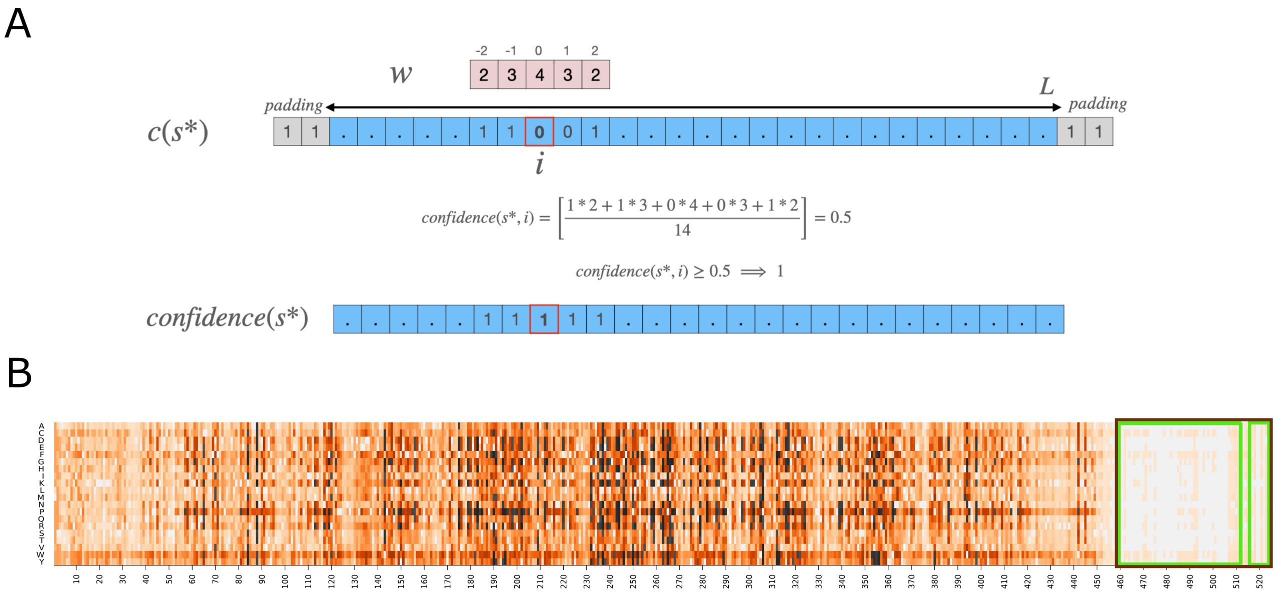


**Figure S24: Overview of the smoothing process for identifying low-confidence residues. A.** The smoothing operation involves performing a convolution over all $c(s^*)$ values in a sequence with a weight vector $W$. The sequence is scanned with overlapping windows and, within each window, the central residue receives the highest weight, while its neighbours receive progressively lower weights. The confidence values $confidence(s^*)$ are obtained by binarising the smoothed scores. **B.** The mutational landscape of the *wdb-PA* (FBpp0084577) protein. The smoothing allows the merging of two nearby low-confidence regions (in green).

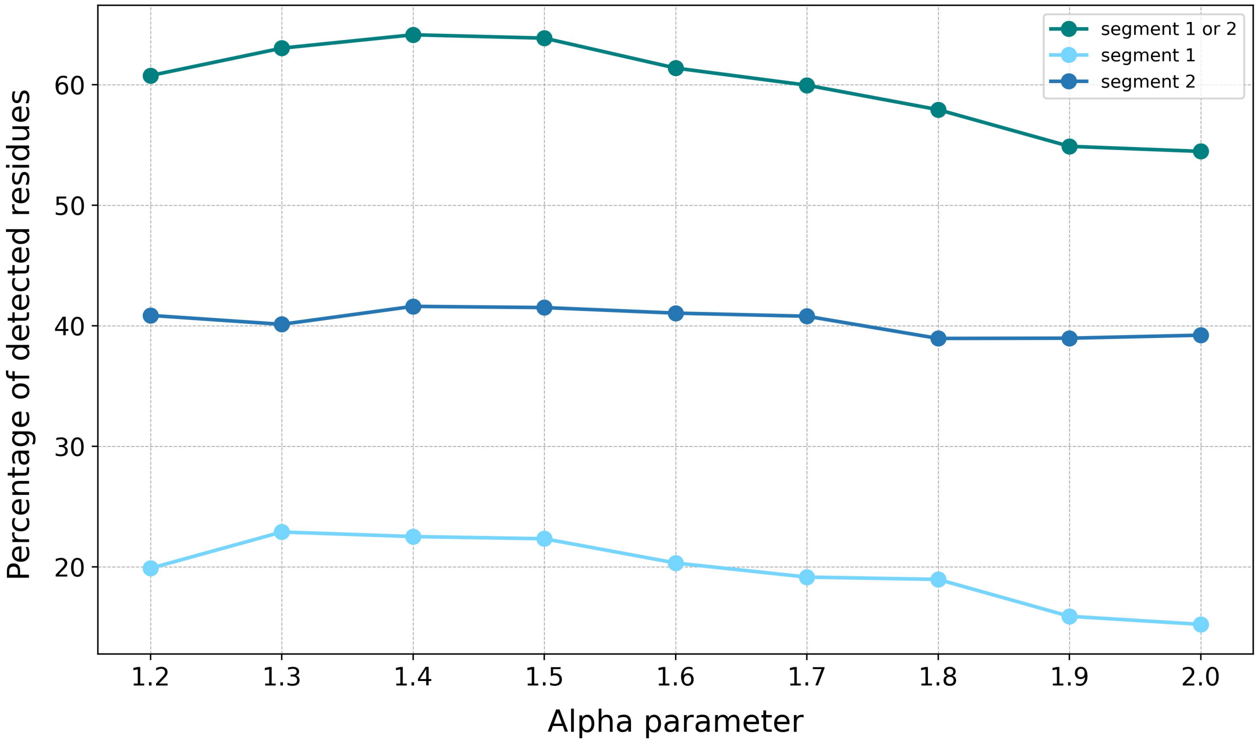


**Figure S25: Assessment of PTM detection in segmented regions as a function of the hyperparameter** $\alpha$. The fraction of detected post-translational modifications (PTMs) within the segmented regions is plotted for segment 1, segment 2, or both. The analysis considers a subset of PTMs from confident proteoforms with an available 3D structure, focusing on regions with pLDDT $< 70$.
